# α-catenin middle- and actin-binding domain unfolding mutants differentially impact epithelial strength and sheet migration

**DOI:** 10.1101/2023.02.01.526618

**Authors:** Jeanne M. Quinn, Yuou Wang, Megan Wood, Annette S. Flozak, Phuong M. Le, Alex Yemelyanov, Patrick W. Oakes, Cara J. Gottardi

**Affiliations:** Department of Pulmonary Medicine, Northwestern University Feinberg School of Medicine, Chicago, IL 60611; Cell & Developmental Biology, Northwestern University Feinberg School of Medicine, Chicago, IL 60611; Department of Cell & Molecular Physiology, Loyola University Chicago Stritch School of Medicine, Maywood, IL 60153

## Abstract

α-catenin (α-cat) displays force-dependent unfolding and binding to actin filaments through direct and indirect means, but features of adherens junction structure and function most vulnerable to loss of these allosteric mechanisms have not been directly compared. By reconstituting an α-cat F-actin-binding domain unfolding mutant known to exhibit enhanced binding to actin (α-cat-H0-FABD^+^) into α-cat knock-out Madin Darby Canine Kidney (MDCK) cells, we show that partial loss of the α-cat catch bond mechanism (via an altered H0 α-helix) leads to stronger epithelial sheet integrity with greater co-localization between the α-cat-H0-FABD^+^ mutant and actin. α-cat-H0-FABD^+^-expressing cells are less efficient at closing scratch-wounds, suggesting reduced capacity for more dynamic cell-cell coordination. Evidence that α-cat-H0-FABD^+^ is equally accessible to the conformationally sensitive α18 antibody epitope as WT α-cat and shows similar vinculin recruitment suggests this mutant engages lower tension cortical actin networks, as its M-domain is not persistently open. Conversely, α-cat-M-domain salt-bridge mutants with persistent recruitment of vinculin and phosphorylated myosin light chain show only intermediate monolayer adhesive strengths, but display less directionally coordinated and thereby slower migration speeds during wound-repair. These data show α-cat M- and FABD-unfolding mutants differentially impact cell-cell cohesion and migration properties, and suggest signals favoring α-cat-cortical actin interaction without persistent M-domain opening may improve epithelial monolayer strength through enhanced coupling to lower tension actin networks.

## INTRODUCTION

α-catenin (α-cat) is a central mechanosensitive scaffold molecule that links the cadherin-catenin complex to the cortical actin cytoskeleton at adherens junctions (AJs), where the latter form structural units essential for tissue morphogenesis and homeostasis (Green et al., 2010; Harris and Tepass, 2010). α-cat mechanosensitivity depends on the conformations and binding activity of its three bundled alpha helical domains (Ishiyama et al., 2013; Rangarajan and Izard, 2012). The N-terminal domain comprises two 4-helical bundles, where the former binds β-catenin (Pokutta and Weis, 2000). The C-terminal domain engages actin through a 5-helical bundle, where force-dependent alteration of the first helix favors high affinity F-actin binding and catch-bond behavior (Buckley et al., 2014; Ishiyama et al., 2018; Wang et al., 2022; Xu et al., 2020). The middle (M)-region of α-cat comprises three 4-helical bundles (M1-3), which undergo sequential unfurling events, M1 at ∼5pN forces followed by M2-M3 at ∼12pN forces, to recruit α-cat M-domain binding partners in a force-dependent manner (Barrick et al., 2018; Kim et al., 2015; Li et al., 2015; Maki et al., 2018; Pang et al., 2019; Seddiki et al., 2018; Terekhova et al., 2019; Thomas et al., 2013; Yao et al., 2014; Yonemura et al., 2010). As such, α-cat is now understood to be a mechanosensitive protein that links acto-myosin force thresholds to distinct α-cat conformational states and partner recruitment. However, the particular adhesive processes and coordinated behaviors that critically depend on α-cat mechanosensitivity are only just emerging (Cho et al., 2022; Donker et al., 2022; Matsuzawa et al., 2018; Monster et al., 2021; Nishimura et al., 2022; Noordstra et al., 2023; Sarpal et al., 2019; Sheppard et al., 2023; Twiss et al., 2012; van den Goor and Miller, 2022).

Our team previously sought to understand the α-cat catch-bond mechanism through a structure-rationalized mutation approach (Ishiyama et al., 2018). By altering 4 amino acids (RAIM to GSGS, a.a. 666-669) in a kinked portion of the first alpha-helix of the α-cat actin-binding domain, Ishiyama et al generated an α-cat with ∼3-fold enhanced F-actin binding *in vitro,* suggesting a model for how acto-myosin forces could alter this region and promote strong actin-binding in cells (Ishiyama et al., 2018). Two recent studies extend and refine this model, implicating the entire first α-helix as a force-gate for F-actin binding, where full loss of this domain (a.a. 666-698) leads to substantially higher actin-binding in vitro (18-fold) (Xu et al., 2020), converting a two-state catch bond into a one-state slip bond (Wang et al., 2022). To accommodate these new findings, we now refer to the Ishiyama et al RAIM → GSGS unfolding mutant as α-cat-H0-FABD^+^, where H0 refers to a short α-helix preceding the first long helix of the α-cat actin-binding domain. This revised nomenclature allows the field to avoid confusion with the α-cat-ΔH1 mutant of Wang et al (2022), which removes both H0 and H1. Since the α-cat-H0-FABD^+^ mutant fails to rescue embryonic lethality of zygotic *αCat* null mutant flies, we know this partial catch-bond mechanism is critical for tissue morphogenesis (Ishiyama et al., 2018). But the specific features of AJ structure and epithelial properties most sensitive to partial loss of α-cat-ABD F-actin catch-bond behavior are less clear, but important in understanding benefits of such regulation for tissue homeostasis. While our previous work showed that expression of α-cat-H0-FABD^+^ in α-cat-negative R2/7-variant DLD1 human colon carcinoma cells leads to epithelial monolayers more resistant to shear-forces, but less efficient at wound repair (Ishiyama et al., 2018), the generality and robustness of these findings remain untested.

In the current study, we generalize these findings to another epithelial cell line, Madin Darby Canine Kidney (MDCK). We demonstrate that partial loss of the α-cat catch bond mechanism (via an H0 α-helix unfolding mutant, RAIM → GSGS) leads to stronger epithelial sheet integrity with greater co-localization between α-cat-H0-FABD^+^ and F-actin, consistent with this mutant’s enhanced association with F-actin under solution-binding centrifugation conditions. However, α-cat-H0-FABD^+^-expressing cells are less efficient at closing scratch-wounds or uniformly packing, suggesting reduced capacity for more dynamic cell-cell coordination. These results imply that cell signals (heretofore unknown) that serve to increase the α-cat/cortical actin interaction might increase epithelial cohesion, but with the cost of limiting cell-cell coordination required for epithelial barrier repair. In addition, while the α-cat/F-actin catch-bond mechanism is thought to be required for M-domain unfolding and partner recruitment, we show that the α-cat-H0-FABD^+^ unfolding mutant is not functionally equivalent to α-cat-M-domain salt-bridge mutants. Indeed, while the latter shows weaker cohesive strength than the former using a standard monolayer fragmentation assay, α-cat-M-domain salt-bridge mutants more strongly perturb the dynamic process of epithelial sheet migration. These data may shed light on how α-cat M-domain missense mutations contribute to epithelial defects that lead to disease (Saksens et al., 2016; Tanner et al., 2021).

## RESULTS

### An α-cat CRISPR knock-out MDCK cell system to interrogate a α-cat mutant function

To more broadly understand how α-cat force-dependent actin binding contributes to epithelial junction organization and function, we sought to generalize previous efforts with a “force-desensitized” α-cat actin-binding mutant (Ishiyama et al., 2018) to the non-cancerous kidney epithelial cell line, MDCK (Madin Darby Canine Kidney), a widely used cell line in the cell-cell adhesion field. We established α-cat knock-out (KO) MDCK cells using the CRISPR-Cas9 system for reconstitution with wild-type or mutant forms of α-cat. We generated RNA guides to α-cat sequences in exons 2 and 4 (Fig. 1A). After transfection and drug selection, single colonies were expanded and screened for lack of α-cat expression. No full length α-cat was detected in knock-out clones by immunoblot analysis (Fig. 1B). Since an immuno-reactive lower molecular weight species was more prominent in clones generated by the most N-terminal guide (gRNA.5), we stably expressed WT α-cat or α-cat-H0-FABD^+^ in the α-cat KO2.2 clone, which expressed this form least. We now know this lower molecular weight species corresponds to a form of α-cat lacking its N-terminal region, likely consequent to our CRISPR-insertion-deletion (INDEL) strategy, which commonly leads to internal ribosome entry and alternative protein products (Tuladhar et al., 2019). As this α-cat isoform lacks the β-cat binding domain and fails to associate with the cadherin-catenin complex, we moved forward with this cell system nonetheless, reasoning this α-cat form cannot directly participate in the cadherin-catenin catch-bond mechanism. Other possible functions of this α-cat isoform expressed at levels seen for guide RNA.5 and generation of complete α-cat null MDCK clones will be described elsewhere (Flozak et al., in preparation). Regardless, the low levels of α-cat variant seen with guide RNA.2 does not obviously localize to cell-cell contacts (Fig. S1A), and restoration of α-cat KO clone 2.2 with α-cat-GFP largely rescues the MDCK parental phenotype and monolayer adhesive strength (Fig. S1B, gray bars).

**Figure 1:**
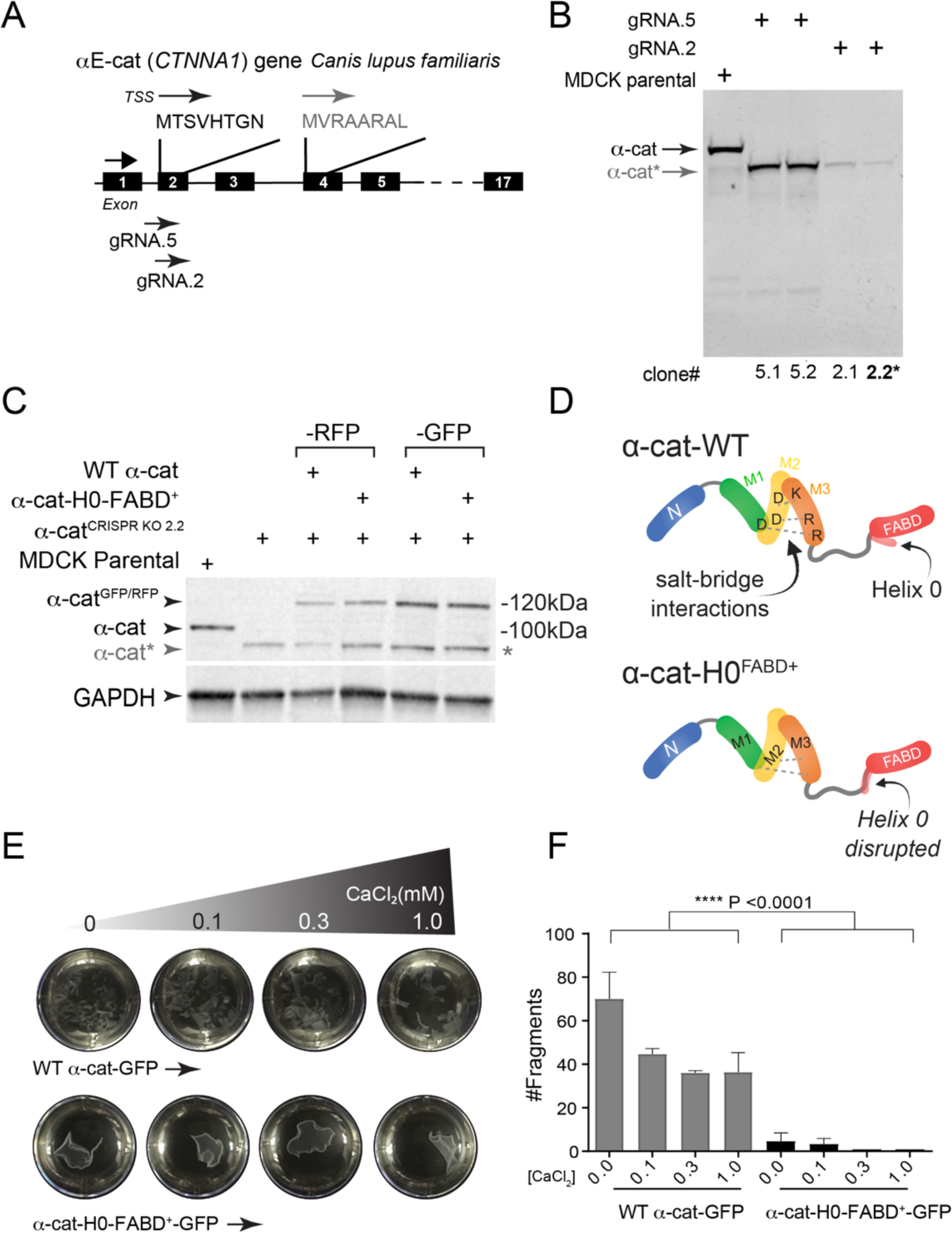
An α-cat actin-binding domain unfolding mutant with enhanced binding to actin filaments increases cell-cell adhesive strength in MDCK cells. (**A**) Generation of α-cat knock-out MDCK cells via CRISPR-Cas9. Schematic shows location of guide RNAs (gRNA.5 and gRNA.2) targeting the MTSVHTGN translational start site (TSS) protein sequence within exon2 of *CTNNA1* gene. (**B**) Immunoblot characterization of clonally expanded α-catenin knock-out lines 5.1, 5.2, 2.1 and 2.2 from gRNA.5 or gRNA2, respectively. Note that gRNA5 led to clones robustly expressing an α-cat fragment (*) likely generated by an alternative translational start site (Bullions et al., 1997). We chose to restore wild-type (WT) α-cat and mutant using α-cat KO clone 2.2. MDCK-parental cell line shown as control (clone 2.2* denotes clone used for reconstitution in this study). (**C**) Immunoblot of MDCK KO2.2 cells transfected with RFP or GFP-tagged versions of WT α-cat or an F-actin-binding-domain (FABD) unfolding mutant (α-cat-H0-FABD^+^). GAPDH was used as loading control. (**D**) Schematic shows location of force-sensitive Helix 0 (H0) in the α-cat actin-binding domain and the H0 unfolding mutant. (**E**) Epithelial sheet disruption assay of MDCK KO2.2 cells expressing WT α-cat (top) or α-cat-H0-FABD^+^ (bottom). Representative monolayers after mechanical stress treatment under a gradient of calcium concentrations (to sensitize calcium-sensitive, cadherin homophilic binding activity) are shown. (**F**) Quantification of monolayer fragments after mechanical disruption using an orbital shaker. Error bars show mean ± SD (n=3, technical replicates), ****p<0.0001, paired t-test.

### An α-cat actin-binding domain unfolding mutant with enhanced actin binding increases epithelial sheet integrity

Flow sorting the brightest third of α-cat-RFP or -GFP expressing cells (not shown), we generated pooled “polyclonal” cell lines to avoid sub-clone variation. Both RFP and GFP-tagged α-cat proteins were expressed similar to normal α-cat levels seen in parental MDCK cells (Fig. 1C-D). Modestly lower expression of RFP-tagged α-cat proteins may be due to expression from different plasmid promoters (Methods). Consistent with evidence that this α-cat-H0-FABD^+^ (RAIM→GSGS) unfolding mutant favors F-actin binding *in vitro* (Ishiyama et al., 2018), we found that MDCK α-cat-KO2.2 cell monolayers reconstituted with α-cat-H0-FABD^+^-GFP remained intact upon mechanical disruption, whereas WT αCat-GFP restored monolayers dispersed into numerous fragments (Fig. 1E-F). These data are consistent with previous work showing that α-cat-H0-FABD^+^ enhances monolayer resistance to mechanical disruption in a human colon cancer cell line (Ishiyama et al., 2018), and generalize the rule that increased α-cat cortical actin interactions can improve the strength of epithelial sheets across tissue types.

### α-cat-H0-FABD^+^ shows enhanced colocalization with F-actin

Restoration of α-cat-KO MDCK cells with a “force-insensitive” mutant that interrupts α-cat Middle-domain salt-bridge interactions (M319G/R326E) led to enhanced recruitment of F-actin and myosin to cell-cell junctions, presumably via α-cat M-domain binding partners that reinforce and amplify acto-myosin contractility (Matsuzawa et al., 2018). Whether a force-desensitized α-cat-H0-FABD^+^ with enhanced actin-binding drives similar actin enrichment at junctions is not known. To address this, α-cat KO2.2 cells restored with WT α-cat-RFP or α-cat-H0-FABD^+^-RFP were grown on porous filters for 10 days before fixing and staining with phalloidin. Curiously, these high-density cultures revealed non-overlapping patches of WT α-cat and F-actin signal, whereas the α-cat-H0-FABD^+^ mutant showed smoother alignment (Fig. 2A). Line-scan analysis of representative bicellular junctions (yellow lines) revealed clear spatial segregation of WTα-cat from F-actin, whereas the α-cat-H0-FABD^+^ mutant distributed more uniformly along F-actin (Fig. 2B). Analysis of full z-stacks confirms greater colocalization of α-cat-H0-FABD^+^ mutant and F-actin compared with WTα-cat (Fig. 2C). Despite the improved colocalization between α-cat-H0-FABD^+^ and F-actin, total actin levels appeared unaltered in WTα-cat-RFP/α-cat-H0-FABD^+^ - GFP co-cultures (Fig. 2D, x-z views, phalloidin in grayscale). Orthogonal views also showed that while WTα-cat-RFP could apically enrich at the zonula adherens junction (Fig. 2D, x-z views, magenta arrows), α-cat-H0-FABD^+^-GFP was distributed more evenly along the lateral membrane, coinciding with F-actin localizations (Fig. 2D, x-z views, green arrows). Together, these data suggest that the α-cat-H0-FABD^+^ mutant likely enhances monolayer cohesion by better engaging actin filaments that run along the entire length of the cell-cell contact. Conversely, the WT α-cat may be tuned to ignore this pool of actin, engaging only with filaments under higher tension thresholds (e.g., at zonula adherens).

**Figure 2:**
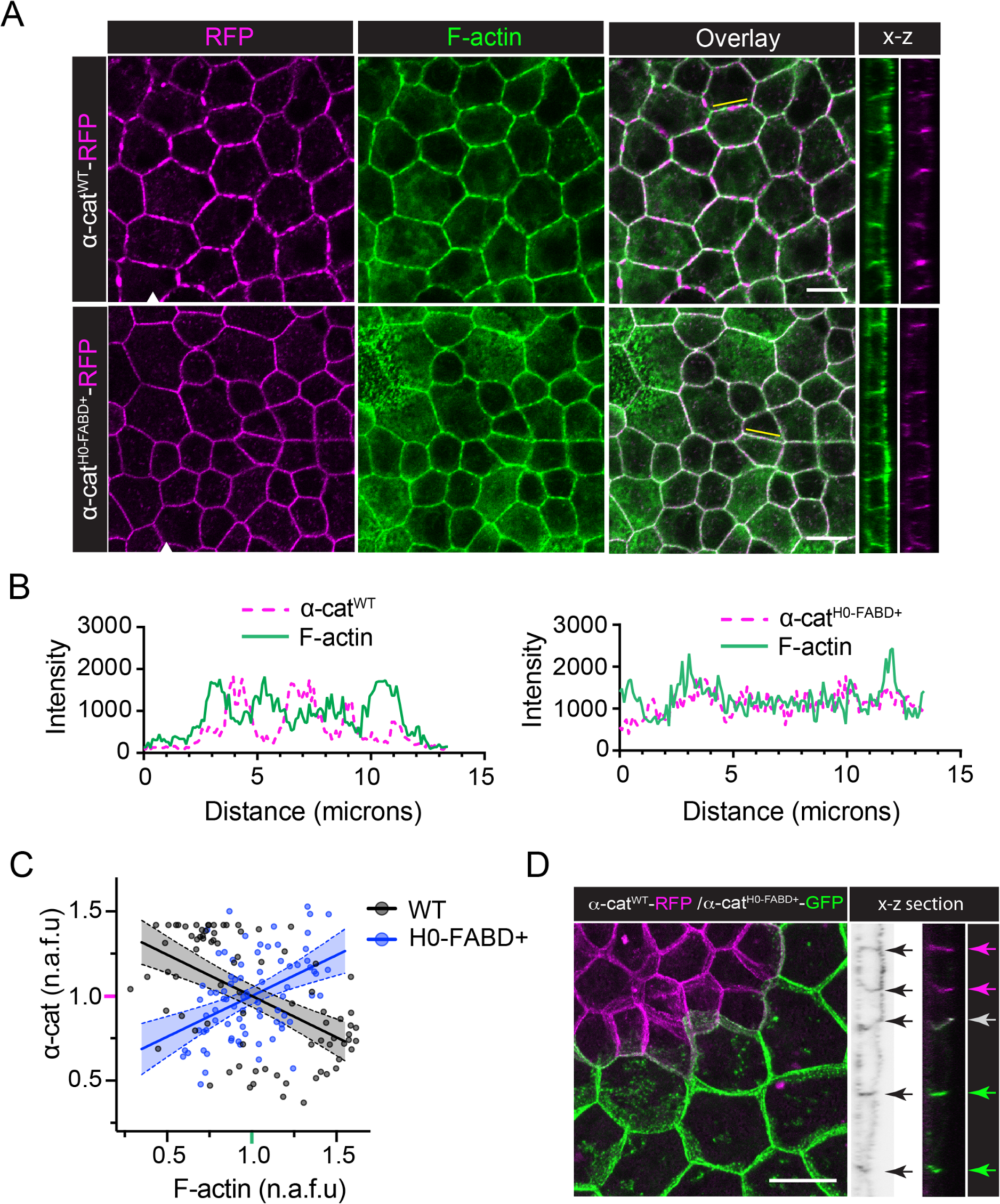
α-cat-H0-FABD^+^ enhances colocalization with F-actin in MDCK cells. **(A)** WT α-cat-RFP and α-cat-H0-FABD^+^-RFP MDCK monolayers (10-12 days on Transwell 0.4μm filters) were stained for phalloidin. Confocal (x-y) and representative x-z orthogonal images (white arrowhead denotes section line) are shown. Bars, 10μm. (**B**) Intensity profiles of α-cat-RFP and F-actin along bicellular junctions (yellow lines in A). (**C**) Colocalization analysis of α-cat-RFP and F-actin (junctions in A). Each symbol represents the normalized intensity (n.a.f.u.) of α-cat and F-actin at 0.1μm junctional ROIs (n=80). Intensity correlation was analyzed by Pearson’s (WT r = -0.536; H0 r = 0.444; ****p < 0.0001) and linear fit plotted with 95% CI (WT slope = -0.488 [-0.662 to -0.315]; H0 slope = 0.482 [0.263 to 0.701]). (**D**) WT α-cat-RFP and α-cat-H0-FABD^+^-GFP were co-cultured and stained for phalloidin (phalloidin shown in grayscale). The inverse correlation between WT α-cat and actin appears most prominent for the C-terminally tagged α-cat lines, which were selected for higher protein expression levels (Fig. S3); we cannot rule-out tag placement as co-contributing to this distribution. Representative x-z orthogonal images are shown. Bar, 10μm. Note α-cat-H0-FABD^+^expression is not associated with elevated levels of F-actin at cell-cell junctions.

### α-cat M1 and M2-domains are not persistently accessible in α-cat-H0-FABD^+^

Single molecule stretch measurements along with steered molecular dynamic simulations reveal how the α-cat M-domain may sequentially unfurl under distinct force thresholds (Leckband and de Rooij, 2014; Li et al., 2015; Yao et al., 2014). In cells, this M-domain unfurled state has been conveniently detected by the monoclonal antibody, α18, which recognizes an M2-helix (a.a. 420-430) preferentially accessible at tension-bearing versus more relaxed cell-cell adhesions (Nagafuchi et al., 1994; Yonemura et al., 2010). Indeed, a previous study revealed the M-domain “activated” salt-bridge mutant (M319G/R326E) was substantially more accessible to α18 than WTα-cat (Matsuzawa et al., 2018). To address the extent to which α-cat-H0-FABD^+^ favors greater accessibility to α18, we co-cultured α-cat-H0-FABD^+^-GFP with WT α-cat-RFP and stained for α18. We find that while α18 can recognize α-cat-H0-FABD^+^ better than WT α-cat in some regions of the monolayer (Fig. 3A), other regions showed preferential recognition of WT α-cat by α18 (Fig. 3B), suggesting α18 accessibility differences between α-cat-H0-FABD+ and WT α-cat are more likely driven by the complex and variable tension forces experienced by the monolayer. We also fail to see constitutive recruitment of vinculin to apical junctions of α-cat-H0-FABD+-expressing cells, where vinculin is known to display tension-dependent recruitment to an α-cat helical region of the M1-domain (Fig. 3C-D) (Hirano et al., 2018; Yao et al., 2014). Collectively, these data reveal that the α-cat M1 and M2-domains are not constitutively accessible in α-cat-H0-FABD^+^, despite its better colocalization with actin and improved monolayer integrity (Figs. 2A and 1E). These data raise the possibility that the α-cat-H0-FABD^+^ mutant engages lower-tension cortical actin networks in dense MDCK monolayers, where these actin networks may be typically ignored by WT α-cat.

**Figure 3:**
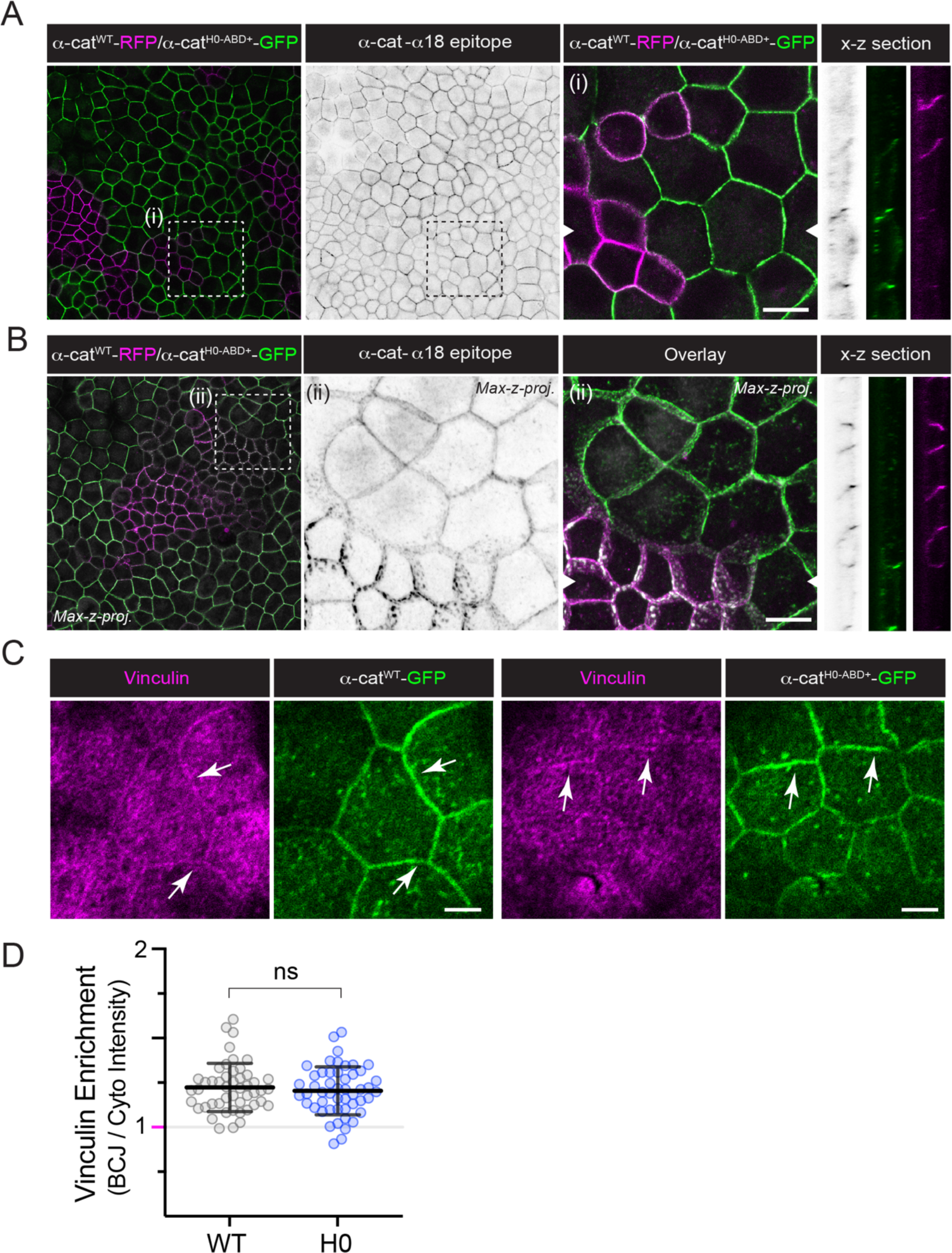
α-cat M2-domain α-18 epitope and vinculin recruitment are similarly accessible to α-cat-H0-FABD^+^ and WT-α-cat. (**A, B**) α-cat M2-domain α-18 epitope shows no preference for α-cat-H0-FABD^+^ over WT-α-cat. WT α-cat-RFP (magenta) and α-cat-H0-FABD^+^-GFP (green) were co-cultured on Transwell filters for 10-12 days, fixed and stained with α18 monoclonal antibody (grayscale). Representative x-z orthogonal images are shown (white arrowhead shows section line). Insets (i) and (ii) show zoomed-in views (right). Bar, 10μm. Note that while the α18 epitope is more accessible in α-cat-H0-FABD^+^ (green) cells in A (x-z grayscale image), B shows the α18 epitope is more accessible in α-cat-WT (magenta) cells. (**C**) Confocal image of filter-grown WT α-cat-GFP and α-cat-H0-FABD^+^-GFP MDCK cells fixed and stained for vinculin (magenta). En face image taken from apical-most plane. Arrows (white) show similarly modest enrichment of vinculin at apical junctions. Bar, 5μm (**D**) Quantification of junctional vinculin enrichment, with each symbol representing the intensity ratio between paired junction (BCJ) and cytoplasm (Cyto) 1μm ROIs (n = 50). Vinculin localization was not affected, as determined by t test p = 0.966.

### α-cat-H0-FABD^+^ interferes with normal cell-cell packing

MDCK cells are often used in epithelial biology for their consistent high-density packing on filters, as evidenced by the normal distribution of apical membrane sizes (Fig. 4A, left). Curiously, MDCK reconstituted with α-cat-H0-FABD^+^ manifest greater heterogeneity in apical membrane size, with small and large sizes often sharing cell-cell contact (Fig.4A, right). Histogram and Kolmogrov-Smirnov analysis reveal α-cat-H0-FABD^+^ alters the distribution of apical areas (Fig. 4B & 4C), despite no significant mean apical areas difference between WT α- and α-cat-H0-FABD^+^ (Fig. 4C). Since this phenomenon can occur in the context of actomyosin contractility differences between adjacent cells (Armon et al., 2018; Matsuzawa et al., 2018), these data suggest an α-cat that is able to engage both lower- and higher-tension actin filaments may be sufficient to propagate signaling relationships that support this irregular cell-packing state.

**Figure 4:**
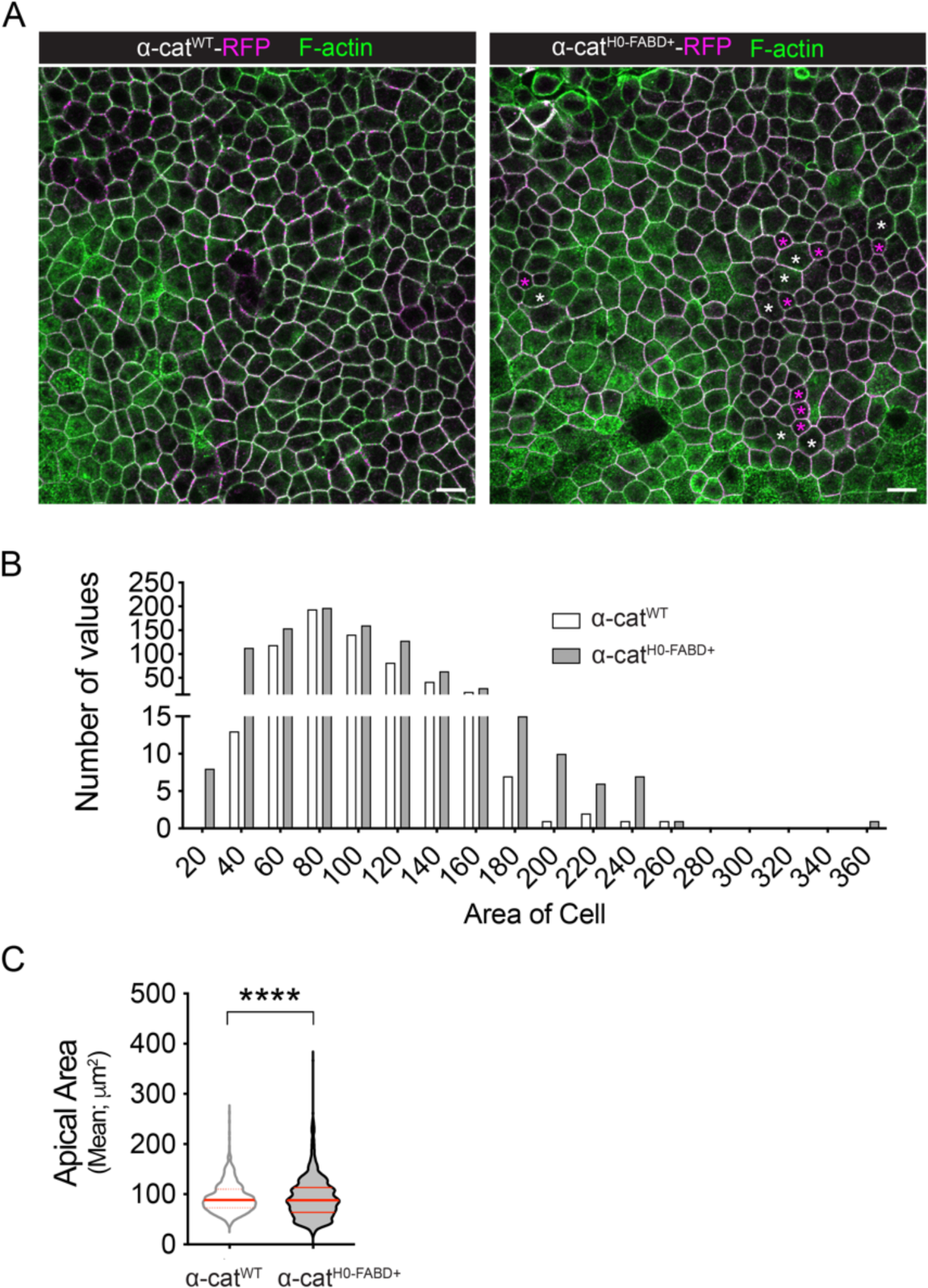
α-cat-H0-FABD^+^ interferes with normal packing of MDCK cells. **(A)** WT α-cat-RFP and α-cat-H0-FABD^+^-RFP-expressing MDCK monolayers (matured 12 days on Transwell 0.4μm filters, magenta) were fixed and stained with phalloidin (green). Magenta asterisk indicates cells with extra small apical area; white asterisk indicates cells with extra-large area. Bar, 10μm. (**B**) Cell number distributions of apical areas in WT α-cat-RFP and α-cat-H0-FABD^+^-RFP cells. Data from three independent FOVs using FIJI-StarDist plug-in. (**C**) Violin plot of apical areas in WT α-cat-RFP and α-cat-H0-FABD^+^-RFP cells. Mean apical areas are not significantly different by unpaired t-test, but the distribution of apical areas (**B**) is significantly different by Kolmogrov-Smirnov analysis (****p < 0.0001). This cell-packing phenotype is strongest for our C-terminally tagged α-cat lines, which also show higher protein expression levels (Fig. S3).

### α-cat-H0-FABD^+^ limits epithelial sheet wound closure

We previously reported that the α-cat-H0-FABD^+^ mutant interferes with wound-front migration using the R2/7 DLD1 cell system (Ishiyama et al., 2018), but whether this also holds for MDCK cells, reflecting a generalized phenomenon is unknown. Live imaging of scratch-wound reveals that α-cat-H0-FABD^+^ mutant cells display unproductive cell-cell tugging orthogonal to the direction of migration (Fig. 5A), leading to reduced wound area closure than WT α-cat (Fig. 5B-C, Movie 1). Finger-like tubular membrane structures, which form via invagination at the interface of leader and follower cells to coordinate directional migration (Hayer et al., 2016), were prominently detected in both WT α-cat and α-cat-H0-FABD^+^ monolayers (Fig. 5A). While some α-cat-H0-FABD^+^ tubular structures appeared less dynamic and more persistent than WT α-cat (Fig. S2A), there was no significant difference in their number or length (Fig. S2B-C, n = ∼50 structures). Interestingly, a greater number of tubular invaginations in α-cat-H0-FABD^+^ wound fronts were angled perpendicularly to the direction migration, consistent with the more uncoordinated (side-to-side) movements of α-cat-H0-FABD^+^ cells relative to WT α-cat (Fig. 5D; S2D-E). These data demonstrate that normal α-cat catch-bond behavior is required for efficient epithelial sheet migration, through favoring a more directional (front-to-back) migration.

**Figure 5:**
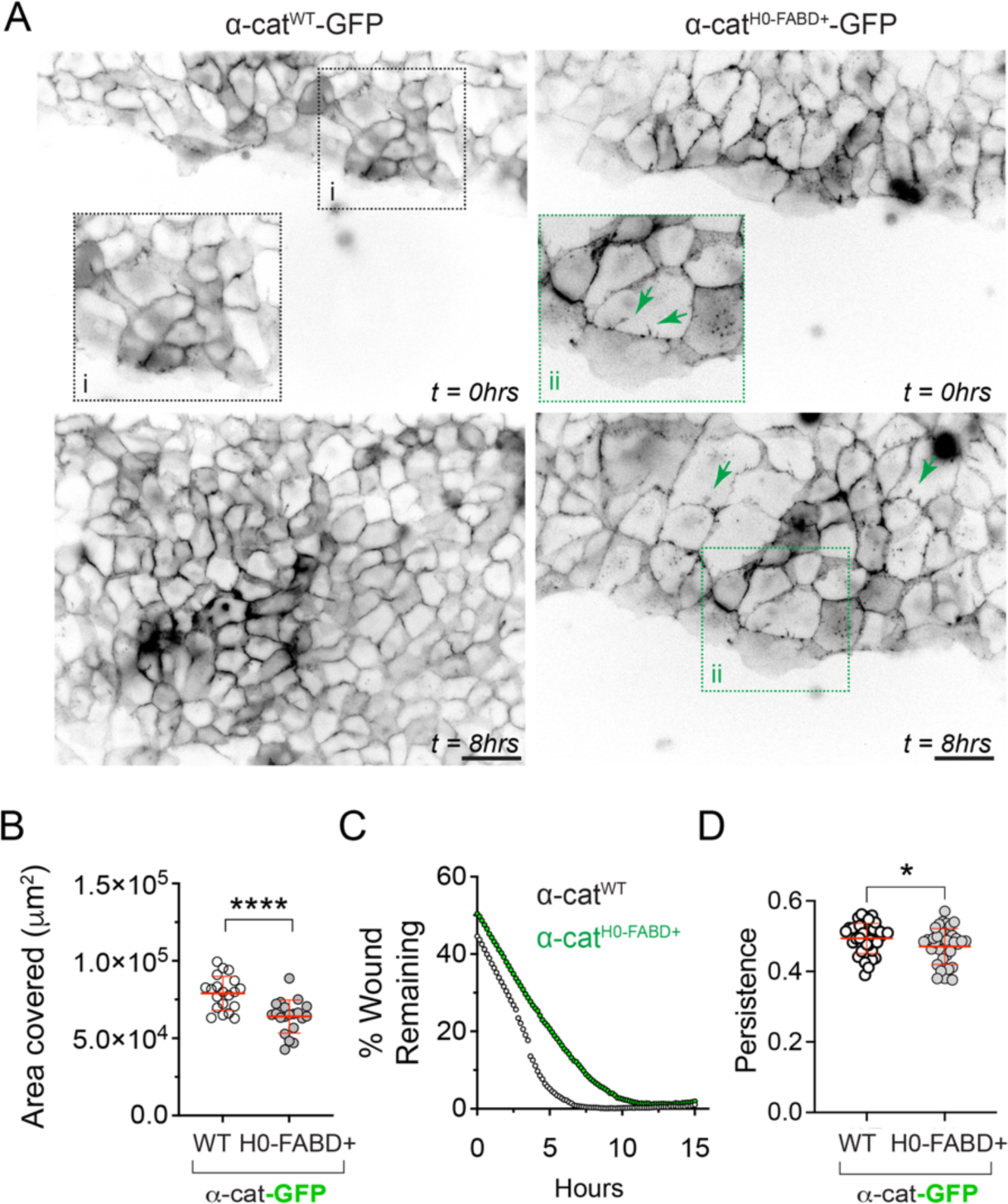
α-cat-H0-FABD^+^ slows MDCK epithelial sheet wound closure. **(A)** Scratch wound healing assays with MDCK KO2.2 cells restored with WT α-cat-GFP or α-cat-H0-FABD^+^-GFP. GFP signals at wound front area at 0-hr or 8-hr are shown. Zoom-in views (i and ii) show “finger-like” protrusions (green arrows). (**B**) Dot distribution plot showing wound closure area covered by WT α-cat-GFP or α-cat-H0-FABD^+^-GFP after 8hrs. Data from three independent repeats, mean ±SD. ****p<0.0001 by ANOVA. (**C**) Representative plot traces showing faster migration rate of WT α-cat-GFP versus α-cat-H0-FABD^+^-GFP cells throughout an entire time-course. (D) Graph showing reduced persistence of α-cat-H0-FABD^+^ cells at wound front compared with WT α-cat cells. Persistence (net displacement/individual cell path length) was quantified using the MTrack FIJI plug-in. Data are presented as mean ±SD with significance by Student t-test * p<0.05.

### α-cat-Middle and F-actin-binding domain unfolding mutants are not equivalent in epithelial monolayer disruption and sheet migration assays

α-cat M-domain “activated” salt-bridge mutants show enhanced recruitment of a number of actin-binding proteins, such as vinculin, afadin and likely others (Matsuzawa et al., 2018; Sakakibara et al., 2020). While destabilization of α-cat M-domain salt-bridges can disrupt epithelial sheet migration (Matsuzawa et al., 2018; Seddiki et al., 2018), the degree to which α-cat M-domain and actin-binding “catch-bond” unfolding mutants are functionally redundant or distinct is not known. To address this question, we compared α-cat M-domain salt-bridge and H0-mutant α-catenins head-to-head using intercellular adhesion and barrier-release assays. Although α-cat constructs with fluorescent-tags placed at the carboxyl-terminus can rescue full development of αCat-null flies (Sarpal et al., 2012), evidence that the unstructured C-terminal extension also contributes to force-activated actin binding (Mei et al., 2020), along with knowledge the extreme C-terminus of α-cat contains a PDZ-interaction motif that might interfere with adherens junction functions, obliged us to generate α-cat mutants tagged with monomeric GFP at the N-terminus. We first affirmed that the N-terminal GFP-tagged α-cat-H0-FABD^+^ migrates more slowly than GFP-α-cat WT cells, similar to α-catenins tagged at the C-terminus (Fig. 6A). While these barrier-release migration differences did not reach statistical significance (p = 0.0664 and 0.0565), they supported scratch-wound data showing an inhibitory role (Fig. 5). The generally faster migration of C-terminally tagged α-catenins may be due to their 2-3 fold higher expression than N-terminally tagged α-catenins (Fig. S3A). Interestingly, our N-terminally tagged unfolding mutants are functionally distinct despite their similar expression by immunoblotting (Fig. S3): While the α-cat H0-mutant shows stronger cohesion than α-cat M-domain salt-bridge mutants (particularly the aspartate to alanine DDD>AAA mutant), the latter is the most potent inhibitor of epithelial sheet migration (Fig 6B). Analysis by particle image velocimetry (PIV) reveals that this due to reduced migratory capacity is associated with less directed and coordinated movements (Fig. 7; Figs S4). Generally, differences in vector speed and direction (Cos theta) were not distinguishable with the first 4-hours of migration, but developed over time with all α-cat mutants significantly differing from WT-α-cat (Fig. 7B-E). Interestingly, α-cat-H0-FABD^+^ monolayers, despite being the most cohesive (Fig. 6C), migrated significantly faster than both α-cat M-domain salt-bridge mutants (KRR and DDD) at later time points (Fig. 7C-D). While α-cat-H0 monolayers showed reduced final speed and correlation across the monolayers compared to WT (Fig. 7E, p = **** by multiple comparisons), H0 monolayers initially demonstrated the largest range of correlated movement compared to WT and M-domain mutants (Fig. 7F, p = **** by multiple comparisons). Spatial correlation increased most in WT monolayers (Fig. 7E-F), coinciding with general velocity and angle trends, suggesting the FABD+ and M-domain mutants do not cohesively ‘respond’ over time like WT monolayers. We noticed that one of our α-cat M-domain salt-bridge mutants (DDD) was particularly potent at inhibiting cell migration (Fig. 7B-E), despite targeting amino acids comprising the KRR salt-bridge pair. While both α-cat M-domain salt-bridge mutants show enhanced recruitment of vinculin, confirming their persistently unfolded state, the DDD mutant showed greater recruitment of vinculin than the KRR mutant (Fig. 8; Fig. S5), offering a possible explanation for different migration characteristics. α-cat M-domain salt-bridge mutants also show increased junctional enrichment in phosphorylated myosin light chain, particularly at tricellular junctions (Fig. 9; Fig. S6, MyoIIA was unchanged, not shown), which may further contribute to less coordinated cellular flows. Together, these data suggest α-cat mutants that favor M-domain unfolding or enhanced actin binding are not equivalent, where M-domain unfolding appears most perturbing to cohesive migration.

**Figure 6:**
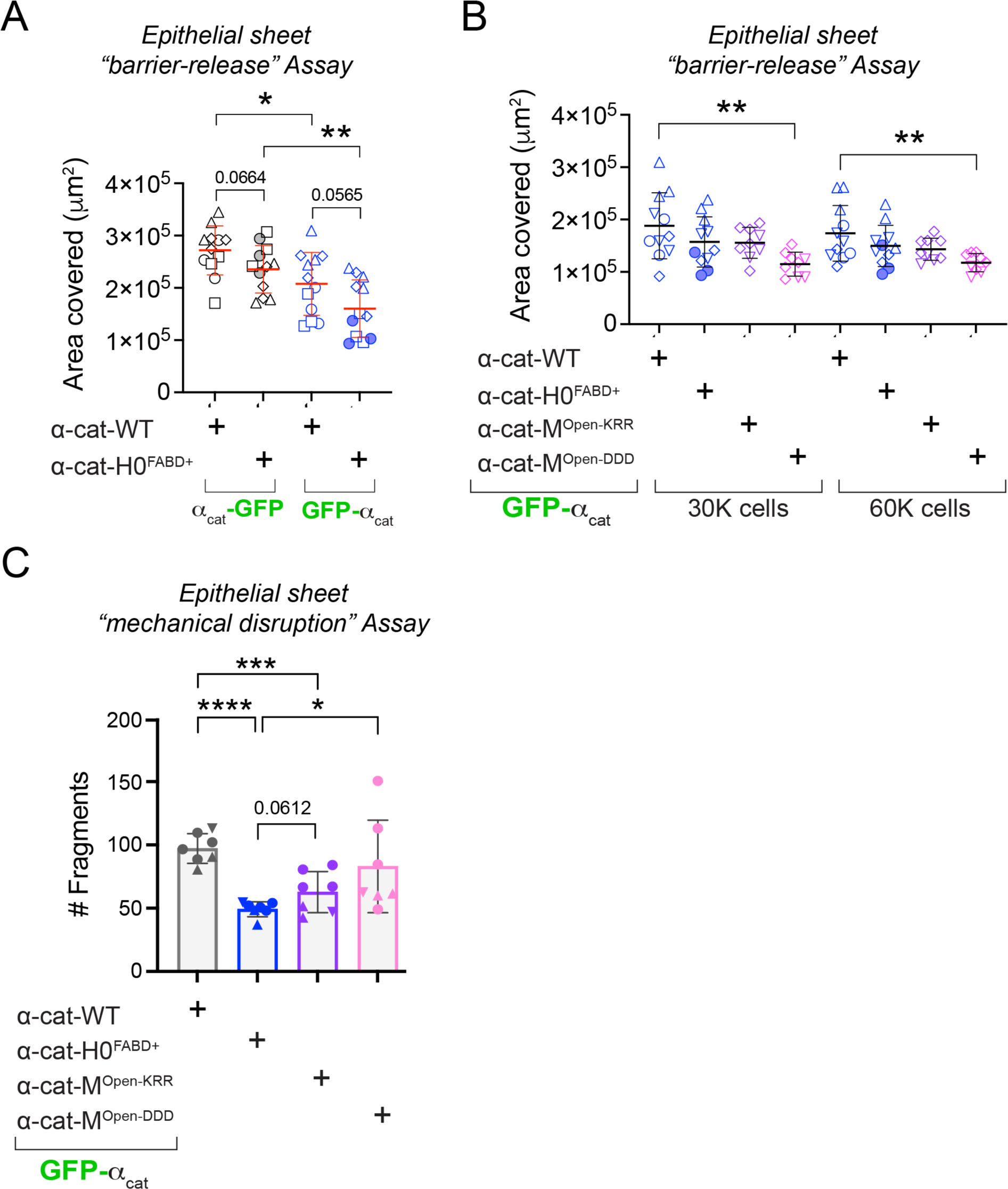
α-cat-Middle domain salt-bridge unfolding mutants perturb epithelial cohesion and sheet migration more than the α-cat-H0-FABD^+^ ‘catch-bond’ mutant. **(A)** Migration capacity of α-cat-WT and α-cat-H0-restored MDCK cells after barrier-removal. Y-axis shows area-covered after 16hrs. N- and C-terminal GFP-tagged versions of α-cat constructs are directly compared. Each symbol reflects a field-of-view quantification. Distinct symbols show biological replicates (n = 4) of barrier-release assays from different days. Note the α-cat-H0-FABD^+^ mutant migrates consistently less than α-cat-WT with p values by ANOVA shown. Data are presented as mean ±SD. Asterisks * p = 0.0218 and ** p = 0.0053. Largest differences in migration rate are due to α-cat expression differences more than GFP-tag placement (immunoblot in Fig. S3A). (**B**) Migration capacity of GFP-tagged α-cat-WT, α-cat-H0-FABD^+^, α-cat-M^OpenKRR^ and α-cat-M^OpenDDD^-restored MDCK cells after barrier-removal. Corresponding immunoblot in Fig. S3A. Each symbol reflects a field-of-view quantification. Distinct symbols show biological replicates (n= 4) of barrier-release assays from different days. Note the α-cat-M^OpenDDD^ mutant migrates significantly slower than α-cat-WT by ANOVA, where ** for 30,000 cell plating p = 0.005 and ** for 60,000 cell plating p = 0.0097). These relative differences in migration do not simply correlate with epithelial sheet strength quantified in (C). (**C**) Monolayer adhesive strength after mechanical disruption. Y-axis quantifies the number of epithelial fragments after Dispase treatment and mechanical shaking. Each symbol reflects measurements from technical replicates (different wells); distinct symbols show biological replicates (n = 3) from assays on a different day with significance by ANOVA, where **** = <0.0001, *** = 0.0007, * = 0.0328. α-cat WT versus α-cat-M^OpenDDD^ = 0.3468 and α-cat-H0-FABD^+^ versus α-cat-M^OpenKRR^ = 0.0612. Data are presented as mean ±SD.

**Figure 7:**
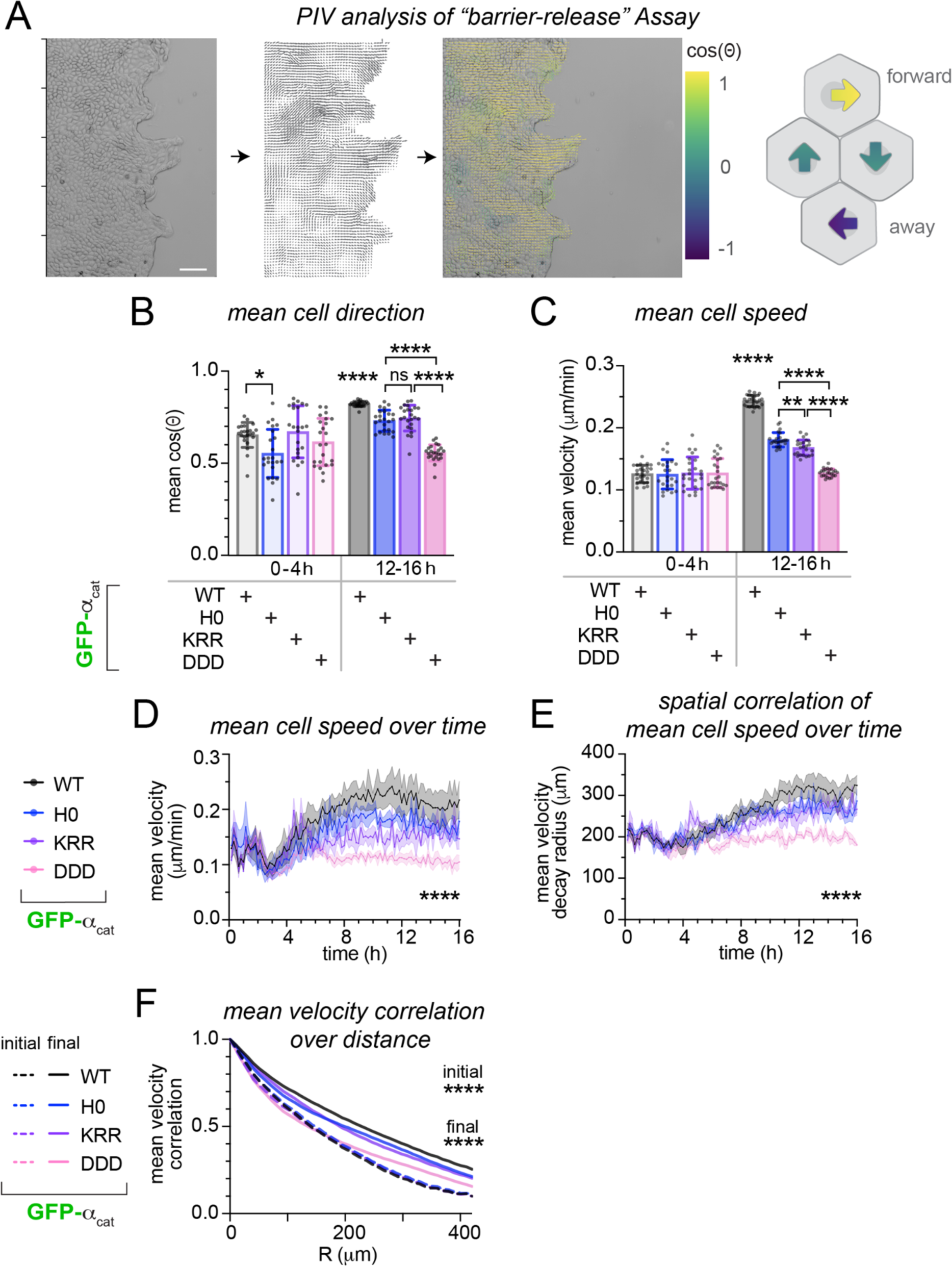
α-cat-Middle domain salt-bridge unfolding mutants perturb collective monolayer speed and coordination more than α-cat-H0-FABD^+^ ‘catch-bond’ mutant. (**A**) Schematic of Particle Image Velocimetry (PIV) analysis of epithelial monolayers. Barrier-release timelapses were background-masked, and calculated flow vectors were analyzed for direction (cos(θ)) and magnitude (velocity). Scale bar = 100μm. (**B**) Average degree of cell-cell coordination during epithelial sheet migration. Y-axis represents the mean cohesive angle, plotted as cos(θ). Data are presented as mean ±SD of 4 biological replicates at initial (t = 0-4 hr) and final (t = 12-16 hr) timepoints, where * = 0.00147, ** = 0.0040, **** = <0.0001 by ANOVA (Dunnett’s multiple comparisons). (**C**) Average velocity of epithelial sheet migration. Data are presented as the mean ±SD of 4 biological replicates at initial and final timepoints, where **** = <0.0001. (**D**) Speed vs. time tracings of monolayers analyzed by PIV. Data are presented as the mean ±SEM of 4 biological replicates with significance by ANOVA, where *** = < 0.0001. (**E**) Spatial correlation of monolayer speed vs time. Data are presented as the mean ±SEM of 4 biological replicates with significance by ANOVA, where **** = < 0.0001. α-cat-WT vs α-cat-H0-FABD^+^ showed no difference in velocity correlation (p = 0.4901), but α-cat-WT showed increased correlation over α-cat-KRR and α-cat-DDD by multiple comparisons (**** = < 0.0001). (**F**) Exponential fit of monolayer velocity correlation as a function of distance (R, μm). α-cat forms show no difference in initial correlation radius, but develop different final correlation radii (*** = 0.0004 by ANOVA); only α-cat-DDD differed by multiple comparisons.

**Figure 8:**
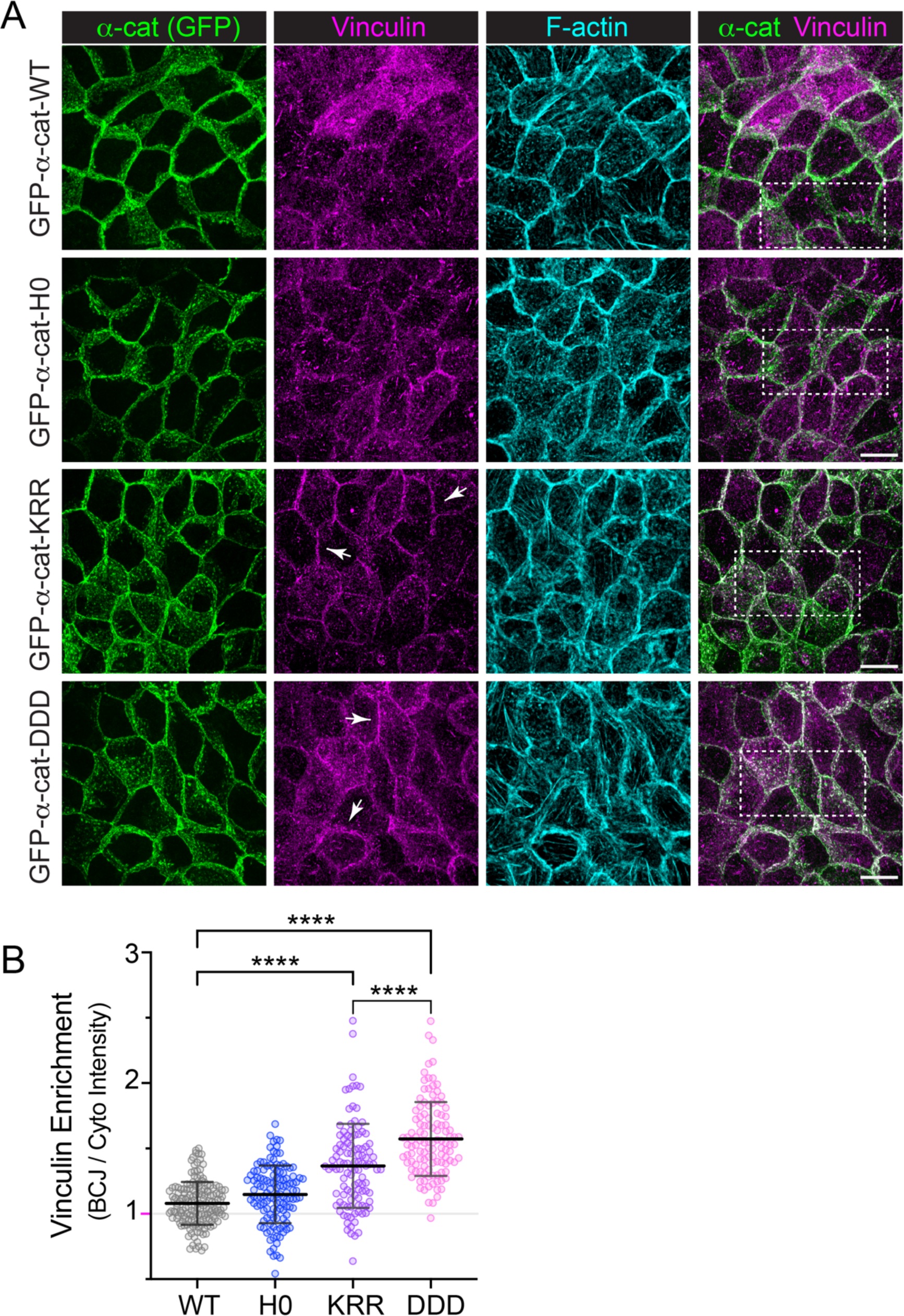
α-cat-M-domain salt-bridge mutants show persistent recruitment of vinculin to cell-cell junctions whereas α-cat-WT and α-cat-H0-FABD^+^ do not. **A**) Immunofluorescence confocal image analysis of α-cat CRISPR-KO 2.2 MDCK cells restored with GFP-α-cat-constructs. Cells grown on glass coverslips were fixed and stained for α-cat (native GFP, green), Vinculin (magenta) and F-actin (cyan). En face images are shown as maximum z-projections. White arrowheads show enhanced recruitment of vinculin to adherens junctions of α-cat-M^OpenKRR^ and α-cat-M^OpenDDD^ constructs relative to GFP-α-cat WT/α-cat-H0-FABD^+^. Overlay image with insets (dotted boxes) are shown at higher resolution in Fig. S6). Scale is 10μm. (**B**) Graph shows junctional vinculin enrichment, with each symbol representing the intensity ratio between paired junction (BCJ) and cytoplasm (Cyto) 1μm ROIs (n = 110+). α-cat constructs affect vinculin localization by ANOVA (**** p < 0.0001) with Turkey’s multiple comparisons (α-cat WT vs H0-FABD^+^ p = 0.0920; α-cat WT vs M^OpenKRR^ and M^OpenDDD^ **** p < 0.0001).

**Figure 9:**
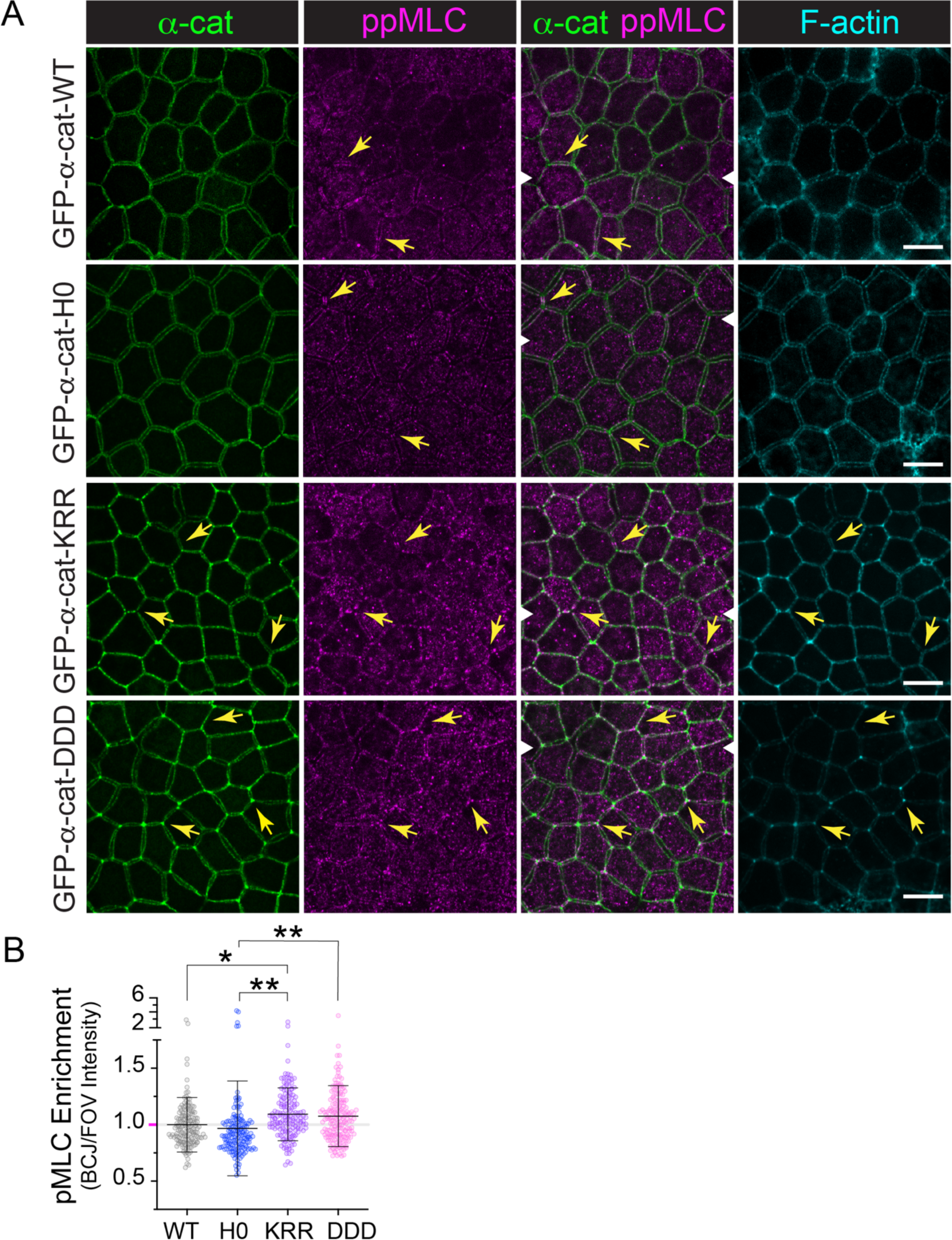
α-cat-M-domain salt-bridge mutants show enrichment of phosphorylated-MLC to cell-cell junctions compared with α-cat-WT and α-cat-H0-FABD^+^. **A)** Immunofluorescence confocal image analysis of α-cat CRISPR-KO 2.2 MDCK cells restored with GFP-α-cat-constructs. Cells grown on Transwell filter supports for two weeks were fixed and stained for α-cat (native GFP, green), ppMLC (magenta) and F-actin (cyan). En face images are shown from a single z-plane. Yellow arrows show enhanced recruitment of ppMLC to adherens junctions of α-cat-M^OpenKRR^ and α-cat-M^OpenDDD^ constructs relative to GFP-α-cat WT/α-cat-H0-FABD^+^. The apparent gap between basolateral membranes, particularly seen in α-cat-WT and -H0-FABD^+^ images, may be due to monolayers being fixed and stored for over a month before staining. Scale is 10μm. White arrowhead in α-cat/ppMLC overlay images shows where x-z section images are taken and shown in Fig. S6. (**B**) Graph shows quantification of ppMLC junctional enrichment, with each symbol representing the intensity ratio of individual bi-cellular junction (BCJ) points 0.1μm ROIs (n = 160-180) ratioed to fluorescence intensity of the entire field of view. α-cat constructs affect ppMLC localization by ANOVA (**** p < 0.0001) with Turkey’s multiple comparisons (α-cat WT vs H0-FABD^+^ p = NS; α-cat WT vs M^OpenKRR^ (*p = 0.03) and α-cat-H0-FABD^+^ vs M^OpenKRR^ or M^OpenDDD^ p**= 0.0012 and 0.0053, respectively).

## Discussion

A number of single molecule studies have shown that α-cat binding to F-actin is force-dependent (Arbore et al., 2022; Buckley et al., 2014; Mei et al., 2020), where the first extended α-helix (H0-H1) of α-cat’s 5-helical bundle actin-binding domain negatively regulates (i.e., force-gates) actin-binding (Ishiyama et al., 2018; Wang et al., 2022; Xu et al., 2020). However, the features of epithelial junction organization and dynamic cell behaviors that critically depend on α-cat/FABD^+^ catch-bond activity remain unclear. Also unclear is the degree to which M- and FABD unfolding mutants manifest distinct consequences for static versus dynamic epithelial adhesive processes. While our previous work showed that restoring α-cat-H0-FABD^+^ in an α-cat-negative colon carcinoma cell line led to epithelial monolayers more resistant to shear-forces, but less efficient at wound repair (Ishiyama et al., 2018), the generality and robustness of these phenotypes remained untested. Here, we extend these findings to the MDCK epithelial cell line. We show that partial loss of the α-cat catch bond property (via an altered H0 α-helix) leads to stronger epithelial sheet integrity with greater co-localization between α-cat-H0-FABD^+^ and F-actin, presumably due to the latter’s persistent association with lower tension cortical actin networks. Since α-cat-H0-FABD^+^-expressing cells are less efficient at closing scratch-wounds or uniformly packing at high density, this mutant displays reduced capacity for dynamic cell-cell coordination. These results reinforce the idea that the α-cat catch bond mechanism is most critical for dynamic rather than static cell-cell adhesions. These results further imply an interesting trade-off; cell signals (heretofore unknown) that serve to reduce the force-sensitivity of the α-cat/cortical actin interaction might improve epithelial monolayer strength through enhanced binding to lower tension cortical actin networks. Indeed, these lower tension actin networks may comprise the bulk of cortical actin in mature epithelial monolayers. But such signals would render α-cat less selective of acto-myosin filaments under higher force, extracting a cost for more dynamic cell migrations, which depend on α-cat ignoring the lower tension actin networks.

A limitation of relying on the α-cat-H0-FABD^+^ mutant to infer α-cat catch-bond activity in cells is the current lack of information on its force-dependent activity using optical trap single molecule analyses (Fig. S7). WT α-cat manifests two-state binding to actin: low or no binding under low forces, binding under mid-forces and slippage under high forces (Arbore et al., 2022; Buckley et al., 2014). Recent studies suggest this two-state model property implicates both H0 and H1, but largely centers on the longer H1 helix (Mei et al., 2020; Wang et al., 2022; Xu et al., 2020). Deletion of both helices (ΔH0-H1 ABD) leads to greatly enhanced actin binding in solution (18-fold; (Xu et al., 2020) and longer binding lifetimes under low forces, but “slippage” at mid-range forces (Wang et al., 2022). As the α-cat-H0-FABD^+^ mutant shows more modest 3-4-fold increase in F-actin binding affinity *in vitro* (Ishiyama et al., 2018) versus 18-fold (Xu et al., 2020), it is likely this mutant still manifests two-state catch bond activity, as the vinculin ABD also lacks H0 yet nonetheless forms catch-bonds with actin (Bakolitsa et al., 2004; Bakolitsa et al., 1999; Borgon et al., 2004; Huang et al., 2017). We reason, therefore, that the α-cat-H0-FABD^+^ mutant has an increased bond lifetime with actin under lower force thresholds in cells. This model is supported by the enhanced colocalization of α-cat-H0-FABD^+^ with lateral membrane F-actin compared with WT α-cat, where the latter accumulates along basolateral membrane patches that exclude actin (Fig. 2). Moreover, evidence M1 and M2-domains are not constitutively accessible to vinculin or the α18 epitope in α-cat-H0-FABD^+^ compared with WT α-cat, further supports the idea that α-cat-H0-FABD^+^ largely engages lower tension cortical actin networks (Fig. 10A, Schematic). Thus, we speculate the α-cat-H0-FABD+ mutant behaves as a “force-desensitized” form of α-cat in epithelial cells, but the particular force-sensitive profile of this mutant and whether it affects the magnitude of the catch-bond behavior will require future testing.

**Figure 10:**
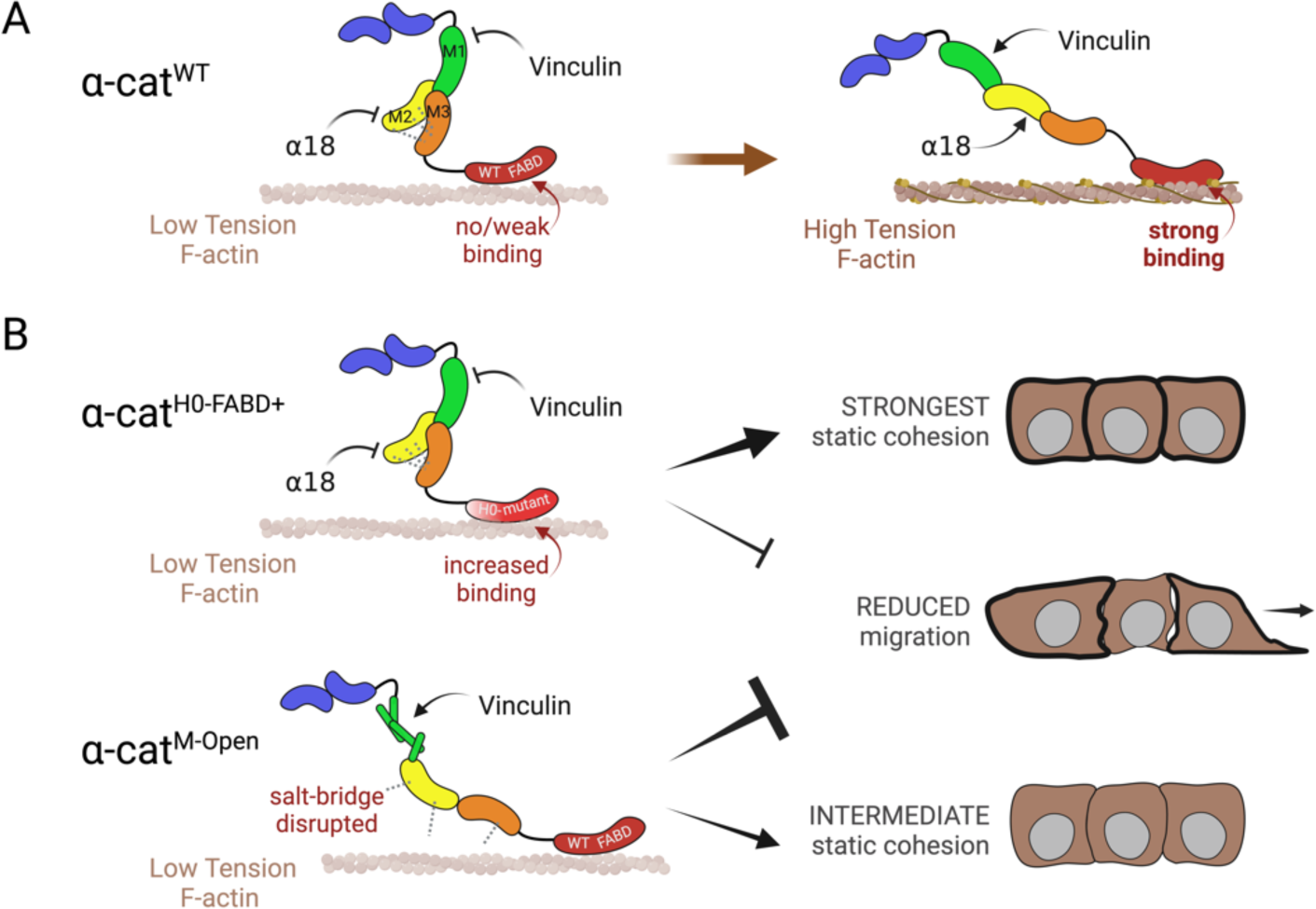
MODEL: Force-desensitized forms of α-cat make distinct contributions to epithelial cohesion, migration and junction organization. (A) Schematic of WT α-cat FABD (red segment of multicolor α-cat protein) binding an actin filament under low tension (left) or high tension (right). “No/weak binding” vs “strong binding” arrows (red) are based on affinity measurements of Buckley et al (2014) and Wang et al (2022) (as in Fig. S7). α-cat N-terminal domains in blue, M1 in green, M2 in yellow and M3 in orange. (B) Schematic of α-cat-H0-FABD^+^ protein binding actin filament under low or no tension (left) based on affinity measurements of Ishiyama et al (2018). Tension-dependent affinity change remains untested for α-cat-H0-FABD^+^ protein, but speculated based on α-cat-ΔH0-H1-FABD^+^ protein analysis of Wang et al (2022) (also in Fig. S7). Similar α18 monoclonal antibody and vinculin accessibility observed for both WT α-cat and α-cat-H0-FABD^+^. Only α-cat-M^OpenKRR^ and α-cat-M^OpenDDD^ show evidence of persistent vinculin and ppMLC recruitment (Fig. 3, 9 and Fig. S5, S6). Activating and inhibitory arrows show consequences of “force-desensitized” α-cat unfolding mutants for epithelial properties: Static epithelial sheet strength is greatest in the α-cat-H0-FABD^+^ mutant, with α-cat-M-domain salt-bridge mutants showing intermediate-to-wild-type levels of cohesion despite increased recruitment of vinculin (and likely other) partners; dynamic epithelial sheet migration is most strongly perturbed in an M-domain unfolding mutant. Created with BioRender.

We were intrigued that cells expressing the α-cat-H0-FABD^+^ mutant led to non-normal packing distribution of MDCK cells grown to high density on filters, where cells with small apically constricted junctions were directly connected to seemingly larger flatter cells (Fig. 4A). While this phenotype may also be related to α-cat expression levels and GFP-tag placement, it is worth noting that tissues can adopt this packing state under different conditions (Armon et al., 2018) and via altered expression of junction proteins and signaling relationships that impact apical contractility (Hildebrand, 2005; Matsuzawa et al., 2018; Sakakibara et al., 2020). Evidence here suggests the possibility that an α-cat with enhanced binding to lower tension actin networks with possible slippage under higher forces may be sufficient to propagate similar cell-cell relationships.

A recent study found MCF7 α-cat KO cells reconstituted with α-cat H0-FABD^+^ showed enhanced adherens junction assembly during the last step of epithelial wound-front closure, where “head-on” junction establishment was associated with reduced lamellipodial activity and cortical actin flow (Noordstra et al., 2023). Interestingly, unliganded surface E-cadherin moved faster in MCF7 cells reconstituted with α-cat H0-FABD^+^, with no change in the overall speed of cortical actin flow. We speculate this enhancement in α-cat H0-FABD^+^ cortical flow may be due to increased binding to lower tension actin networks, which at nascent cell-cell contacts allows for a greater number of actin-engaged cadherin-catenin complexes to drive adherens junction formation.

Lastly, α-cat is long known to bind F-actin directly or indirectly through force-dependent recruitment of other actin-binding proteins, such as vinculin, but the degree to which these modes of actin-binding are redundant or distinct has not been directly tested. By comparing α-cat H0-FABD^+^ and α-cat M-domain salt-bridge mutants head-to-head in epithelial sheet disruption and barrier-release migration assays, we show that M-domain unfurling is more disruptive to adhesion and migration than enhancement of F-actin binding. This finding is notable and may be related to evidence that familial missense mutations that cause Macular Pattern Eye Dystrophy exclusively localize to the M-domain (Saksens et al., 2016; Tanner et al., 2021), underscoring the importance of this domain to normal α-cat function. Future work will be required to understand the force-sensitive targets of α-cat M-domain unfolding or dysfunction that negatively impact coordinated epithelial cell behaviors (Fig. 10B, Schematic).

## ACKNOWLEDGEMENTS

This work relied on the following Northwestern University services and core facilities: Flow Cytometry (NCI CA060553); Center for Advanced Microscopy (NCI CCSG P30 CA060553) and Skin Biology and Diseases Resource-based Center (P30AR075049) from the NIAMS. We thank Pankaj Bhalla for gene editing advice with help from the SBDRC GET iN Core.

## Author contributions

JMQ, MNW, ASF, PML and AY conducted experiments and analyzed results; PWO advised and wrote code for the PIV analysis; CJG designed and supervised the study. JMQ, YW, PWO and CJG wrote the manuscript. PWO and CJG provided funding for project.

## METHODS

### Plasmid generation and constructs

The αE-catenin cDNA was amplified by PCR and subcloned into either a modified pcDNA3.1 vector (Ishiyama et al., 2010) to express αE-catenin-mRFP or a modified pCAN vector (Ichii and Takeichi, 2007) to express αE-catenin-mEGFP (EGFP with the dimerization-disrupting A206K mutation (Zacharias, 2002). N-terminally tagged αE-catenins were synthesized by VectorBuilder using a dimerization-disrupted mEGFP (A206K) in third-generation lentiviral vectors with components pLV[Exp]-CMV>mEGFP-αE-catenin EF1A(core)>Puro. Lentivirus packaging (psPAX2, #12260) and envelope (pMD2.G, #12259) plasmids were purchased from Addgene.

### Cell culture and stable cell line selection

MDCK II cells were maintained in Dulbecco’s Modified Eagle’s Medium (DMEM, Corning), containing 10% fetal bovine serum (FBS, Atlanta Biologicals or JRS Scientific), 100 U/ml penicillin and 100 μg/ml streptomycin (Corning). α-cat/*Ctnna1* knockout MDCK cells were generated using CRISPR-Cas9 system, RNA guides were designed targeting α-cat sequences in exons 2 and 4. Guide RNA (gRNA) targeting different canine *CTNNA1* exons were designed with CHOPCHOP online tools (Labun et al., 2019). Sequences of oligonucleotides were as follows: 5’-GAAAATGACTTCTGTCCACACAGG-3’ (exon 2, covering the MTSVHTG protein sequence) and 5’-AGTCTAGAGATCCGAACTCTGG-3’ (exon 2, covering the SLEIRTLA sequence). Single guide RNA (sgRNA), Cas9 nuclease (HiFi) and duplex buffer were purchased from Integrated DNA Technologies. RNAs were reconstituted and diluted to 5 μM with duplex buffer; Cas9 protein to 10 μM with PBS. MDCK cells were plated in 12-well plates at a seeding density of 1.0 x 10^5^ cells in 1 mL DMEM-complete a day before. For one reaction, sgRNA (20 μL), Cas9 (15 μL), DMEM (30 μL, no serum and antibiotics) and Lipofectamine RNAiMax (4 μL, Invitrogen) were mixed and incubated at room temperature for 20 min. The complex was treated to the cells in complete medium and incubated overnight at 37 °C and 5% CO2. The medium was changed and the cells were allowed to recover for one day. Cells were split to maintain 30-50% confluency. The sgRNA-Cas9 treatment was repeated 3 times. Cells were expanded and sorted with a flow cytometer (FACSMelody, BD) to 96-well plates to grow single cell colonies. The colonies were screened for low (< 5 ng/well at confluent) α-cat expression using a α-cat C-terminal antibodies. The selected colonies were verified with western blot. Knockout cell line (2.2 clone) was chosen based on lowest level of α-cat isoform lacking N-terminal sequences.

After transfection and drug selection, colonies were expanded and screened for lack of α-cat expression. Cells were transfected with expression vectors by using Lipofectamine 3000 (ThermoFisher Scientific), and cells with stable protein expression was selected based on antibiotic resistance (100 μg/mL Zeocin for pcDNA3.1 and 300 μg/mL hygromycin for pCAH vectors) and subsequently isolated by flow cytometry. For lentivirus production, 293T cells (GeneHunter) were transfected with 8ug expression vector (Vector Builder), 6μg psPAX2, and 2μg pMD2.G using TransIT (Mirus). Viral supernatant was collected 48 and 72h after transfection, passed through a 0.45μm filter, and supplemented with 1μL/mL polybrene (Sigma). To generate stable GFP-α-cat lines, MDCK α-cat KO cells were transduced for 6hr at 37°C on 10cm plates with 2mL prepared viral supernatant. Cells were selected in culture media containing 5μg/mL puromycin, then sort-matched for GFP using a FACSMelody 3-laser sorter (BD).

### Antibodies

The following primary antibodies were used: polyclonal rabbit anti-α-cat (C3236, Cell Signaling), hybridoma mouse anti-α-catenin (5B11, (Daugherty et al., 2014)), Rat anti-α-catenin mAb (α18) was a kind gift of Dr. A. Nagafuchi (Nara Medical University, Nara, Japan), rabbit anti-GAPDH (Cat# sc-25778, Santa Cruz, now discontinued), polyclonal rabbit anti-GFP (A11122, Invitrogen) and Phalloidin-488 or -568 (A12379, Invitrogen). Secondary antibodies for Western blotting included HRP-conjugated goat anti-mouse and -rabbit antibodies (Bio-Rad), or fluorescently labeled donkey anti-mouse and -rabbit antibodies (680RD or 800RD, LiCor Biosciences). Secondary antibodies for immunofluorescence included IgG Alexa Fluor 488 or 568-conjugated goat anti-mouse or -rabbit antibodies (Invitrogen).

### Immunofluorescence and Imaging

Cells were grown on cell culture inserts (Falcon), fixed in 4% paraformaldehyde (Electron Microscopy Services, Hatfield, PA) for 15’, quenched with glycine, permeabilized with 0.3% Triton X-100 (Sigma), and blocked with normal goat serum (Sigma). Primary and secondary antibody incubations were performed at RT for 1h, interspaced by multiple washes in PBS, and followed by mounting coverslips in ProLong Gold fixative (Life Technologies). Images of α-cat, F-actin, and vinculin localizations were captured with Nikon A1R Confocal Laser Point Scanning microscope using NIS Elements software (Nikon) with GaAsP detectors and equipped with 95B prime Photometrics camera, Plan-Apochromat 60x/1.4 objective. Confocal Z-stacks were taken at step size of 0.3μm.

### Image Analysis and Fluorescence Quantification

To examine the localization of vinculin and α-cat, junctional enrichment was quantified on single apical slices in FIJI and normalized. Briefly, the integrated signal intensity from 1μm circular ROIs were measured from bicellular junctions (BCJ) and the adjacent cytoplasm (Cyto). Cyto signal was subtracted from BCJ signal, and the enrichment was normalized to 1 within each experimental group. Junctional phospho-myosin was similarly quantified, but enrichment was taken over FOV signal to minimize the contribution of background cytosolic antibody signal. To compare the distribution of F-actin and α-cat along junctions, 0.1μm circular ROIs were placed along BCJs and the integrated signal intensity was measured from both channels. The junctional signal was standardized to the FOV signal and normalized to 1. Normalized F-actin vs. α-cat signal for each ROI was plotted, and correlation was measured with Pearson’s “R” value and by linear fit. F-actin vs α-cat signal is further reported with a typical line-scan analysis taken along a BCJ indicated in the figure panel.

### Scratch Wound Assay

MDCK cells (250,000) were plated for 24hr on LabTek #1 4-well chamber slide (43300-776, Thermo Scientific), wounded with a P200 micropipette tip and allowed to recover for 2hr. Prior to imaging, DMEM media was replaced with FluoroBright DMEM (Life Technologies) and 10 μg/mL Mitomycin C (Sigma) to limit cell proliferation. Cells in Figure 5 were imaged with the 20x objective every 10 min (both phase contrast and fluorescent channels) on the Nikon Biostation IM-Q with the slide holder module (located in Nikon Imaging Facility) at 37°C, 5% CO2, for 15 hr. 6 fields of view (FOV) were captured along the wound edge. To quantify change in wound area, the resulting .ids file was imported to ImageJ, and the wound edge of the phase-contrast image was traced with the polygon tool at time = 0 and time = 15hr. The area of the resulting polygon was measured in pixels² and the resulting data were compared by performing One-way ANOVA statistical analysis followed by Tukey’s multiple comparison test using GraphPad. Images presented in the paper were adjusted for brightness/contrast but were otherwise unprocessed. Individual cell tracks were visualized using the MTrackJ plug-in (FIJI) (Fig. S2E). Cadherin fingers were counted manually and within 150-minutes after wounding. Length, intensity & angles were measured using FIJI by using the line tool (line width of 3 pixels) along the cadherin fingers at 200% zoom.

### Epithelial barrier-release assay

Cells (30,000 and 60,000) were plated to 35mm 4-well Ibidi plates (#80466) at for 48h. Two hours before imaging, the barrier was pulled, and media was changed to DMEM-10%FBS with 10μg/ml mitomycin C for 2 hours incubation. Media was changed to Fluorobrite media with 10% FBS. Positions were selected along the wound edge and brightfield images were acquired at 10-minute intervals over 16h hours on a Nikon Ti2 microscope using a 20x air objective. The monolayers’ coordinated movement was further analyzed by Particle Image Velocimetry (PIV) on background-masked timelapses.

### Particle Image Velocimetry (PIV) and Flow Analysis

To analyze the collective motion of migrating cells during barrier release assays, we performed particle image velocimetry (PIV) measurements using code from openpiv (Liberzon et al., 2021). Briefly, the original image was broken down into a series of small windows that covered the entire image. The displacement of each window between successive frames was determined by performing a cross-correlation. This resulted in a vector field encoding the local movement of objects in the image. For this analysis we used a window size of 90×90 pixels (33×33 µm) and a slightly larger search window size of 93 pixels. Each window overlapped by 65 pixels (∼24 µm), resulting in a grid that produced a displacement vector every 28 pixels (∼10 µm– approximately the size of a single cell). Once the vector field was calculated they were first masked to only include displacements in regions where there were cells, and filtered to remove outliers. All vectors outside of the masked region were set to zero. Vectors that had a magnitude of greater than 30 pixels/frame (1.1 µm/min) or a signal-to-noise correlation ratio of less than 1.035 were considered outliers. These vectors were removed and replaced with a vector representing the local mean of the surrounding vectors with a kernel size of 3 pixels.

Once displacement fields were measured, we performed additional analyses to determine the average magnitude and direction of the velocity vectors in each frame over time. The average direction was calculated by determining the cosine of the angle between the displacement vector and a unit vector that points along the x-axis (i.e. towards the open area, cos(8)). These values were averaged over all vectors in the masked region of the image. To calculate the local correlation of these values within the field, we performed an autocorrelation of the masked vector field with itself (Angelini et al., 2010). The resulting correlation was normalized to 1 and then radially averaged. A correlation length was determined by fitting this curve to a single exponential decay *C*(*r*) = e^−*r/R*0^, where *R*_0_ represents the decay length. All analyses were performed in python. Code and example data are available at https://github.com/OakesLab.

### Epithelial sheet-disruption Assay

Epithelia resistance to mechanical disruption was measured using a “Dispase assay” as previously described (McEwen et al., 2014; Wood et al., 2017). Cells (400,000) were plated on 12-well plates (Corning) for 48hr, then washed and pre-incubated for 15mins with calcium-free HBSS + 20mM HEPES. Cell monolayers were lifted from plate wells by incubating at 37°C with a filtered solution of HBSS, 20mM HEPES, 20mg/mL dispase II (Roche), 0.3μM calcium, and 0.5μM magnesium. Monolayers were subjected to a shaking force of 1,400 rpm (Thermomixer R, Eppendorf) in 1min increments. Each well was imaged with an iPhone 13 using the 0.5X lens before and after shaking, and the resulting images were analyzed as TIFs in FIJI. Image files were blindly counted for macroscopic fragments using the BlindAnalysisTools plugin, taking the average count from duplicate photos of each well, with 200 serving as the upper limit for fragment counting.

## Key resources Table

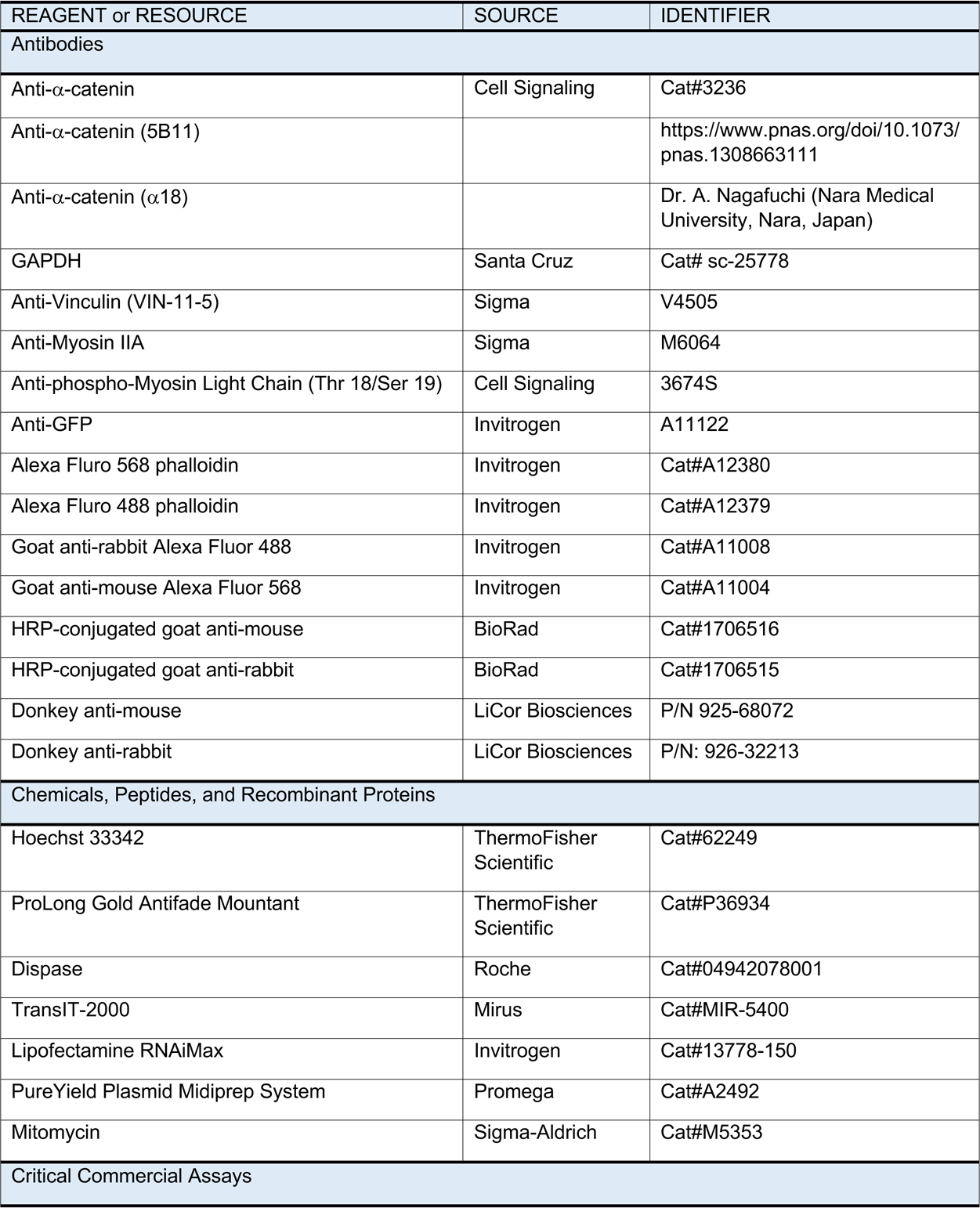
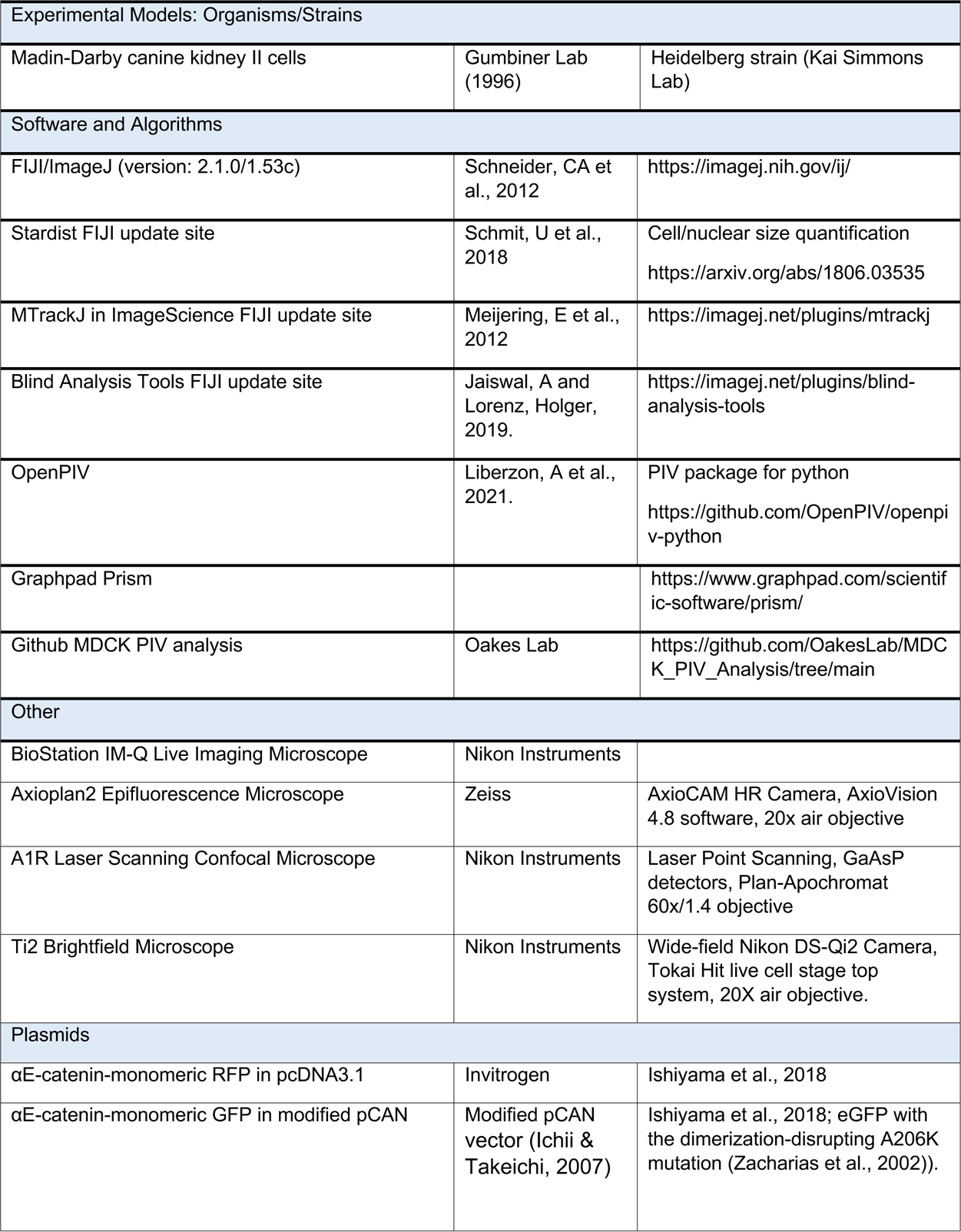
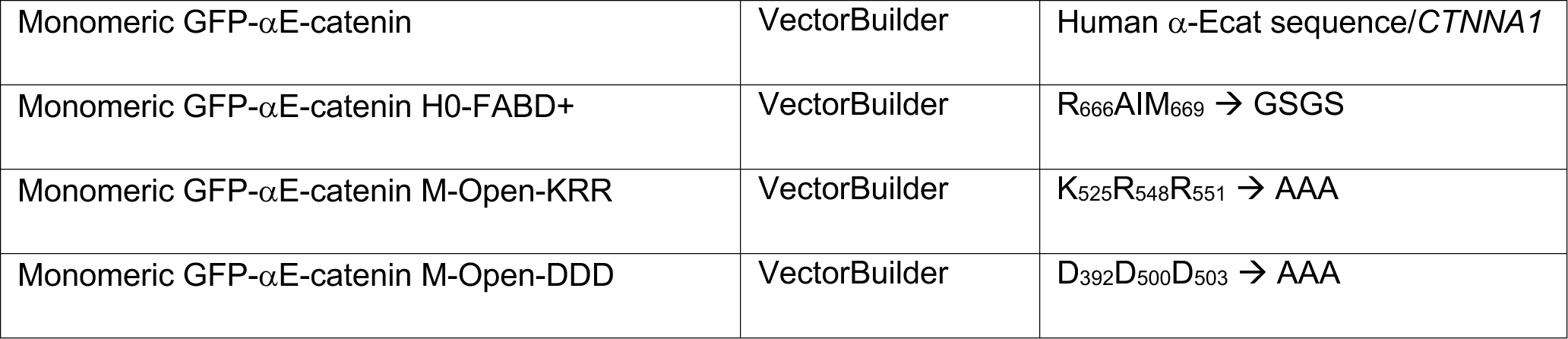

---

**Figure S1:**
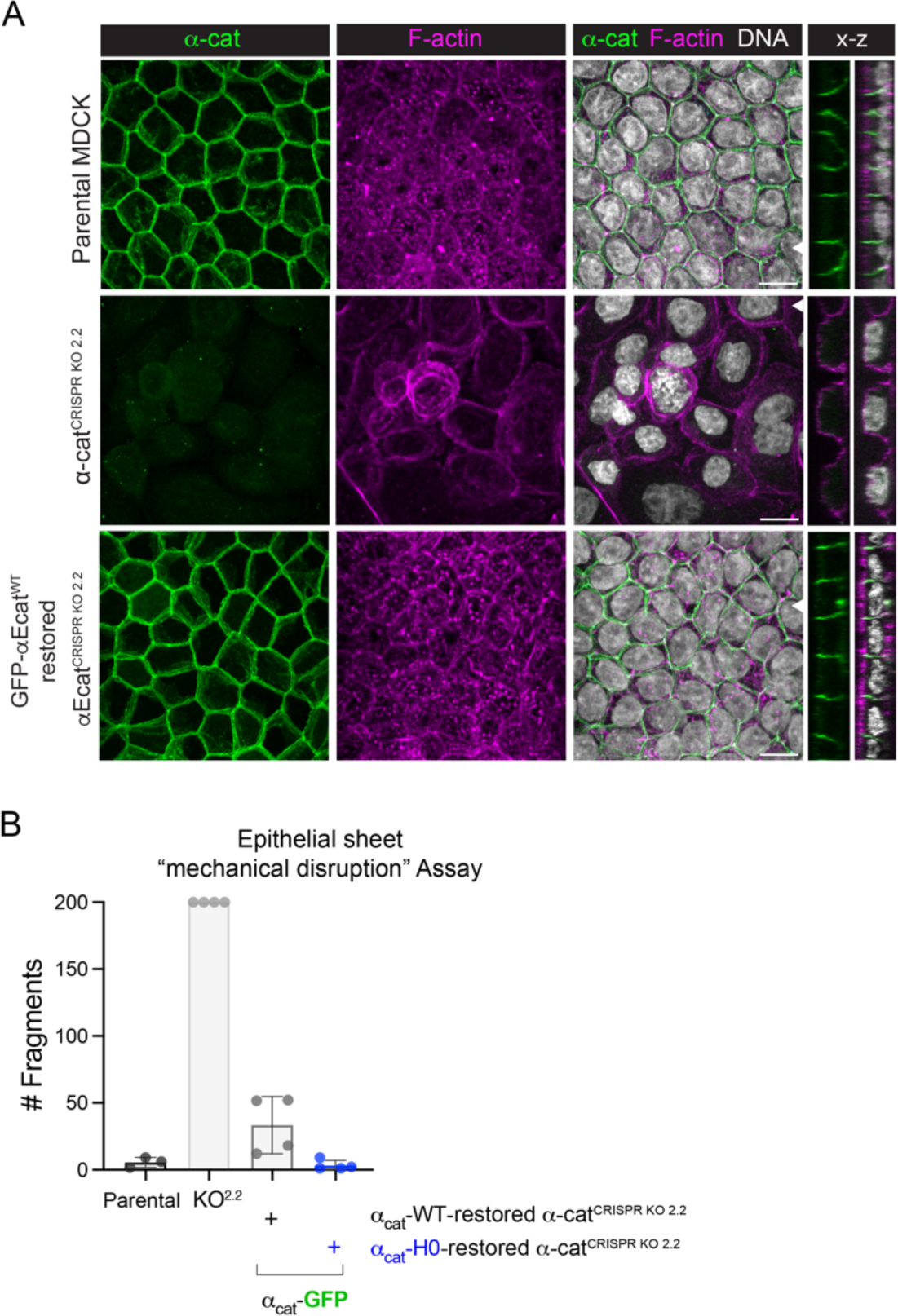
Characterization of α-cat CRISPR-KO 2.2 MDCK cells. (**A**) Immunofluorescence confocal image analysis of parental MDCK cells (top row), α-cat CRISPR-KO 2.2 MDCK cells (middle row) and GFP-α-cat-restored α-cat CRISPR-KO 2.2 MDCK cells (bottom row) stained for α-cat (green), F-actin (magenta) or DNA (gray). En face images are shown as maximum z-projections with x-z optical sections to right. White arrowhead shows localization of optical z-section. Scale is 10μm. Note α-cat short isoform expression in CRISPR-KO 2.2 MDCK cells (Fig. 1B) shows no clear localization to cell-cell contacts. (**B**) Graph shows quantification of MDCK monolayer adhesive strength after Dispase-treatment and mechanical shaking (as in Fig. 1E-F). Note wild-type α-cat-GFP-restored CRISPR-KO 2.2 MDCK cells rescue monolayer integrity to a similar extent as parental MDCK cells.

**Figure S2:**
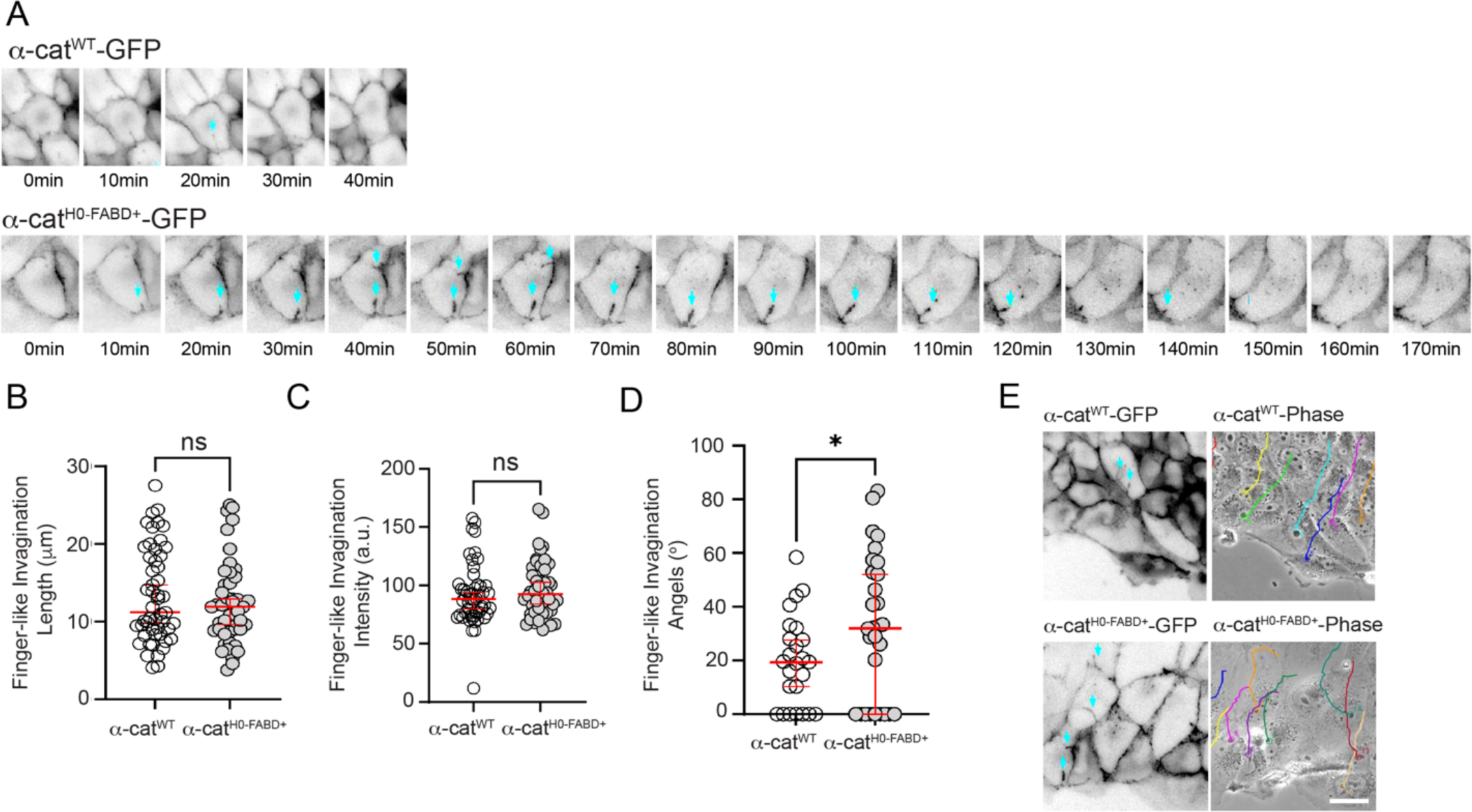
Finger-like tubular invaginations form equally between WT α-cat and α-cat-H0-FABD^+^ expressing MDCK wound front leader and follower cells. *Supplemental images to* Figure 5 **(A)** Time-lapse images of MDCK KO2.2 cells restored with WT α-cat-GFP or α-cat-H0-FABD^+^-GFP. Native GFP fluorescence shown in grayscale (inverted). Cyan arrows point to finger-like invaginations at interface between leader cell and follower cell. (**B-D**) Quantification of tubular invaginations (n = 51 for WT α-cat and 46 for α-cat-H0-FABD^+^: (**B**) Length, unpaired t-test, p = 0.4101; (**C**) Fluorescence intensity, unpaired t-test, p = 0.3477; (**D**) Direction/angle formed by tubular invagination and direction of sheet migration, Kolmogorov-Smirnov test, where asterisk (*) reflects p = 0.0408. (**E**) Representative WT-α-cat tubular invaginations (cyan arrows) align with direction of migration (phase image, colored tracks); α-cat-H0-FABD^+^ invaginations align perpendicular to direction of migration. Scale bar, 5μm.

**Figure S3:**
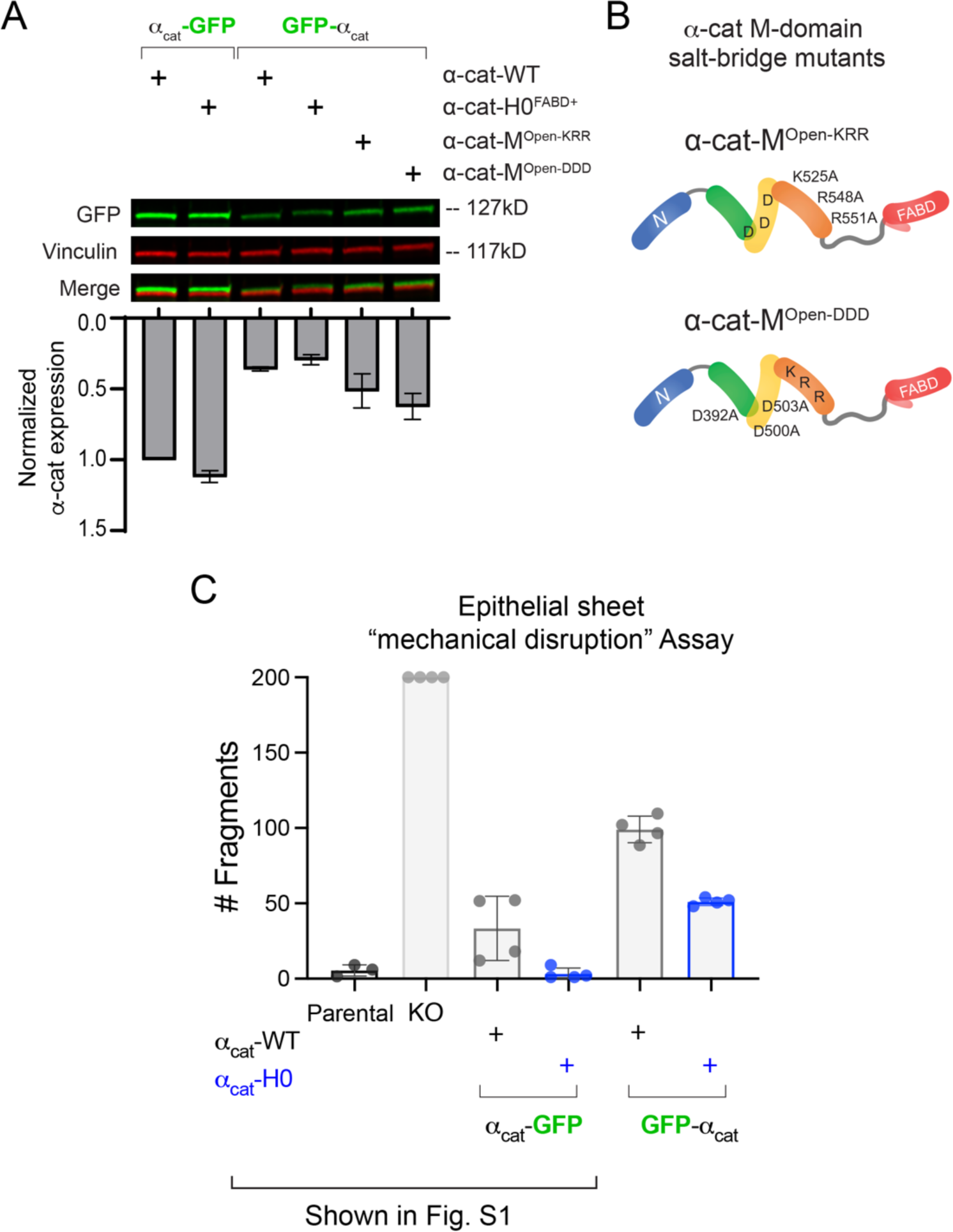
Characterization of GFP-tagged α-cat cell lines. (**A**) Immunoblot of N- and C-terminal GFP-tagged α-cat constructs. Vinculin (red) is used as a loading control. Graph shows α-cat protein expression normalized to Vinculin. Data are presented as mean ±SD (n = 3). Higher protein expression of C-terminally tagged α-cat relative to N-terminally tagged α-cat is due to different flow cytometry gating strategies used to select these cell lines. Modest differences in protein expression between GFP-α-cat WT/α-cat-H0-FABD^+^ versus α-cat-M^Open^ constructs are evident, but do not explain differences in epithelial sheet strength or migration seen in Fig. 6. (**B**) Schematic of α-cat-M^OpenKRR^ and α-cat-M^OpenDDD^ constructs, showing location of salt-bridge mutant modified residues. (**C**) Epithelial monolayer adhesive strength assay after mechanical disruption. Y-axis quantifies the number of epithelial fragments remaining after Dispase treatment and mechanical shaking. Each symbol reflects measurements from technical replicates (different wells). The first portion of this graph was shown in Fig. S1. Here we show cohesive strength of N- versus C-terminally tagged α-cat cell lines. Note that parental MDCK cells show strongest intercellular adhesion compared with α-cat CRISPR-KO2.2 cells restored with GFP-tagged-α-catenins. Cells expressing higher levels of α-cat are less prone to fragmentation (see immunoblot in A).

**Figure S4:**
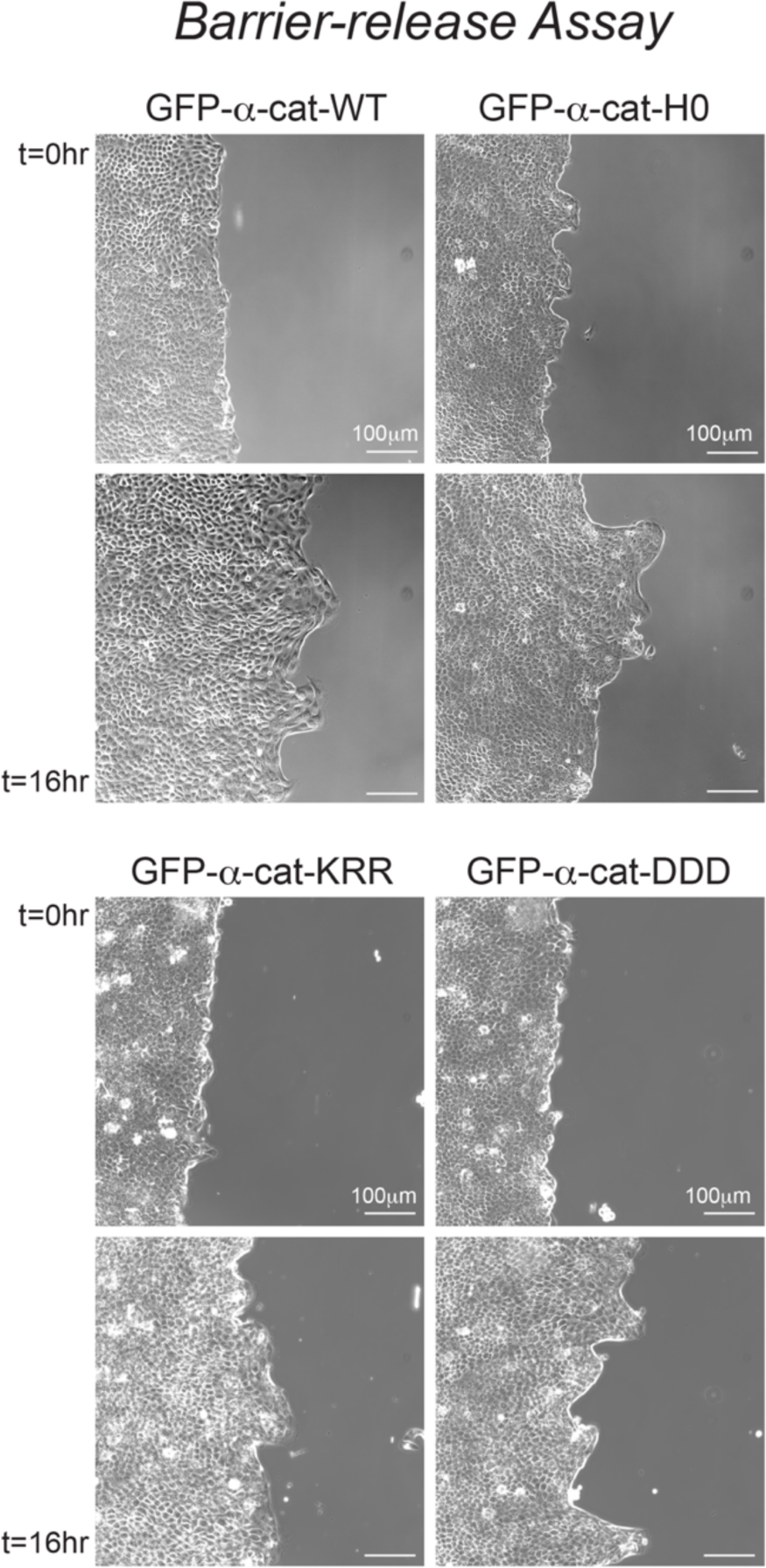
Barrier-release epithelial sheet migration assay. *Supplemental images to* Figure 7. Phase images of GFP-α-cat-restored MDCK cells plated in 35mm 4-chambers Ibidi plates (60,000 cells per well). Two hours before imaging, the barrier was pulled, and media was changed to DMEM-10%FBS with 10μg/ml mitomycin C for 2 hours incubation. Media was changed to Fluorobrite media with 10%FBS. Positions were selected along the wound edge and brightfield images were acquired using 20x objective at 10-minute intervals over 16h hours on Ti2 microscope (Nikon). Scale bar = 100μm. Specific cell lines/α-cat constructs are indicated.

**Figure S5:**
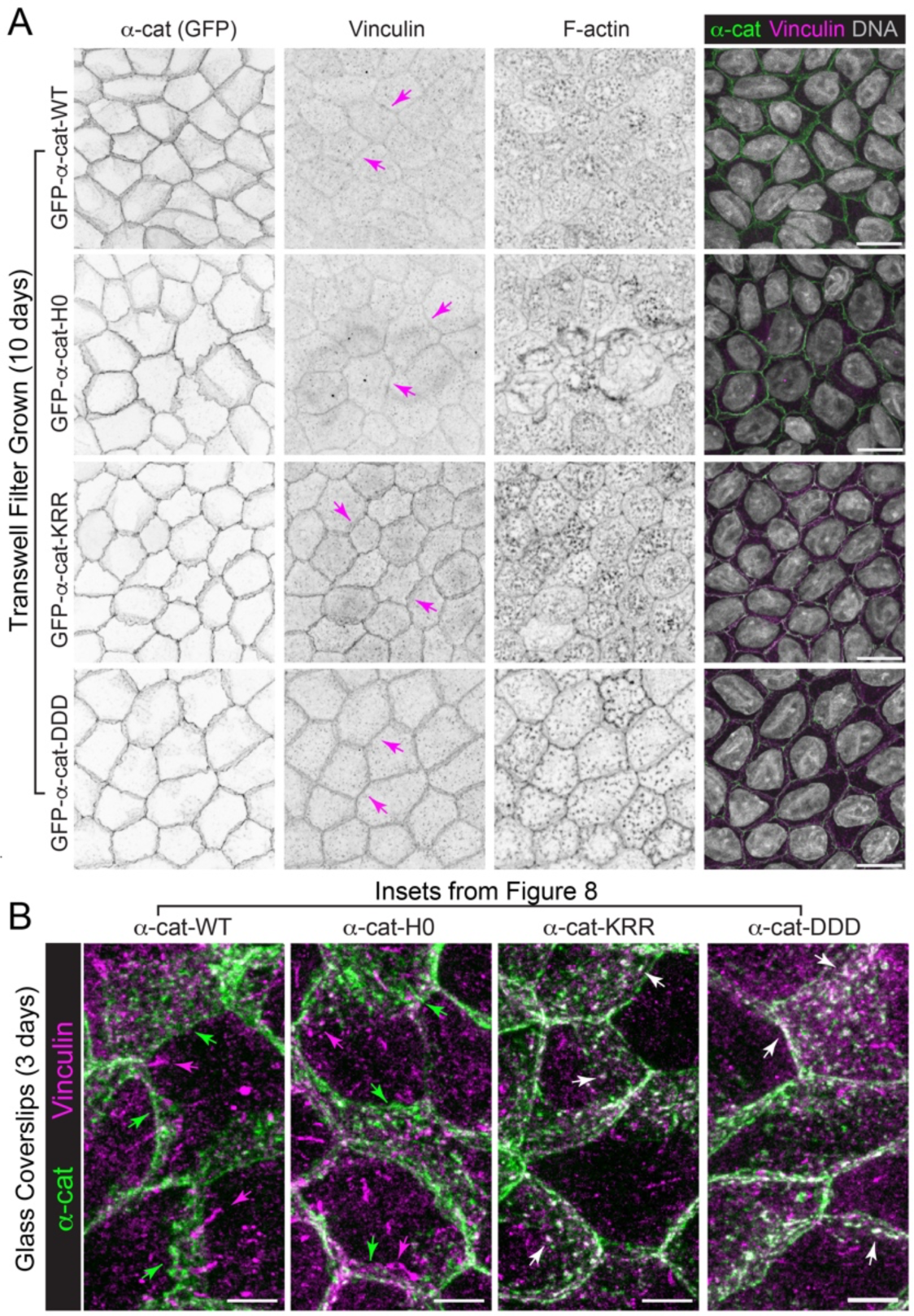
α-cat-M-domain salt-bridge mutants show persistent recruitment of vinculin to cell-cell junctions compared with α-cat-WT and α-cat-H0-FABD^+^. *Supplemental images to* Figure 8. A) Immunofluorescence confocal image analysis of α-cat CRISPR-KO 2.2 MDCK cells restored with GFP-α-cat-constructs. Cells grown on Transwell filter supports for two weeks were fixed and stained for α-cat (native GFP, green), Vinculin (magenta) and F-actin (cyan) (overlay). Individual en face images are shown as maximum z-projections (inverted grayscale). Magenta arrows show enhanced recruitment of vinculin to adherens junctions of α-cat-M^OpenKRR^ and α-cat-M^OpenDDD^ constructs relative to GFP-α-cat WT/α-cat-H0-FABD^+^. Overlay image with nuclei (gray). Scale is 10μm. (**B**) Inset images from Fig. 8 (dotted boxes) show evidence of α-cat clusters along lateral membranes (cells grown on glass coverslips). Note α-cat cluster size is not obviously different, but co-localization with Vinculin is more clearly evident in α-cat M-domain salt-bridge mutants (white arrows). Scale is 5μm.

**Figure S6:**
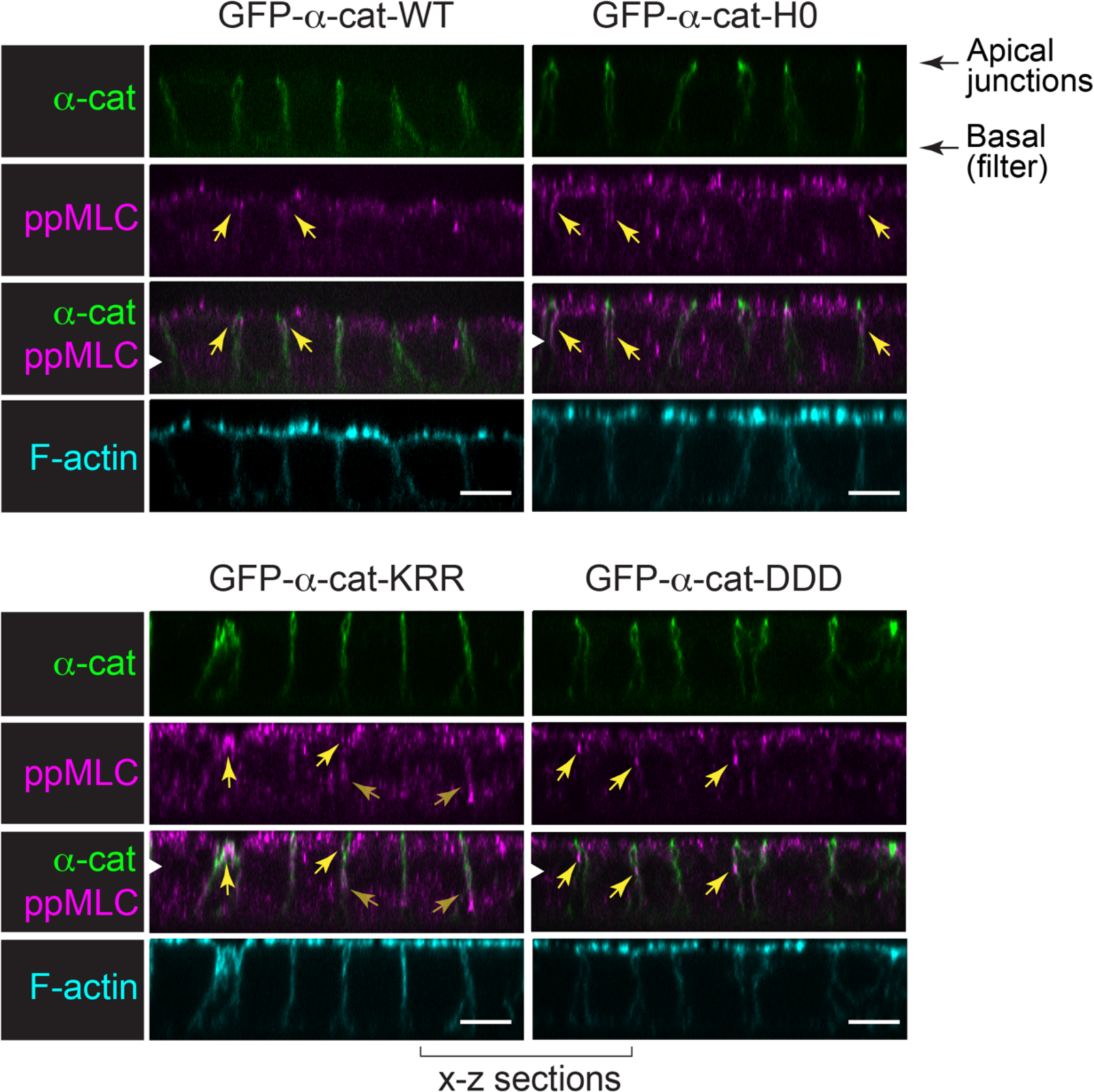
α-cat-M-domain salt-bridge mutants show enhanced recruitment of phospho-Myosin Light Chain (ppMLC) to adherens junctions compared with α-cat-WT and α-cat-H0-FABD^+^. *Supplemental images to* Figure 9. Immunofluorescence confocal image analysis (x-z views) of α-cat CRISPR-KO 2.2 MDCK cells restored with GFP-α-cat-constructs. Cells grown on Transwell filter supports for 1-2 weeks were fixed and stained for α-cat (native GFP, green), ppMLC (magenta) and F-actin (cyan) (overlay). Yellow arrows show enhanced recruitment of ppMLC to adherens junctions of α-cat-M^OpenKRR^ and α-cat-M^OpenDDD^ constructs relative to GFP-α-cat WT/α-cat-H0-FABD^+^. White arrowheads show z-plane corresponding to x-y images in Figure 9. Scale is 10μm.

**Figure S7:**
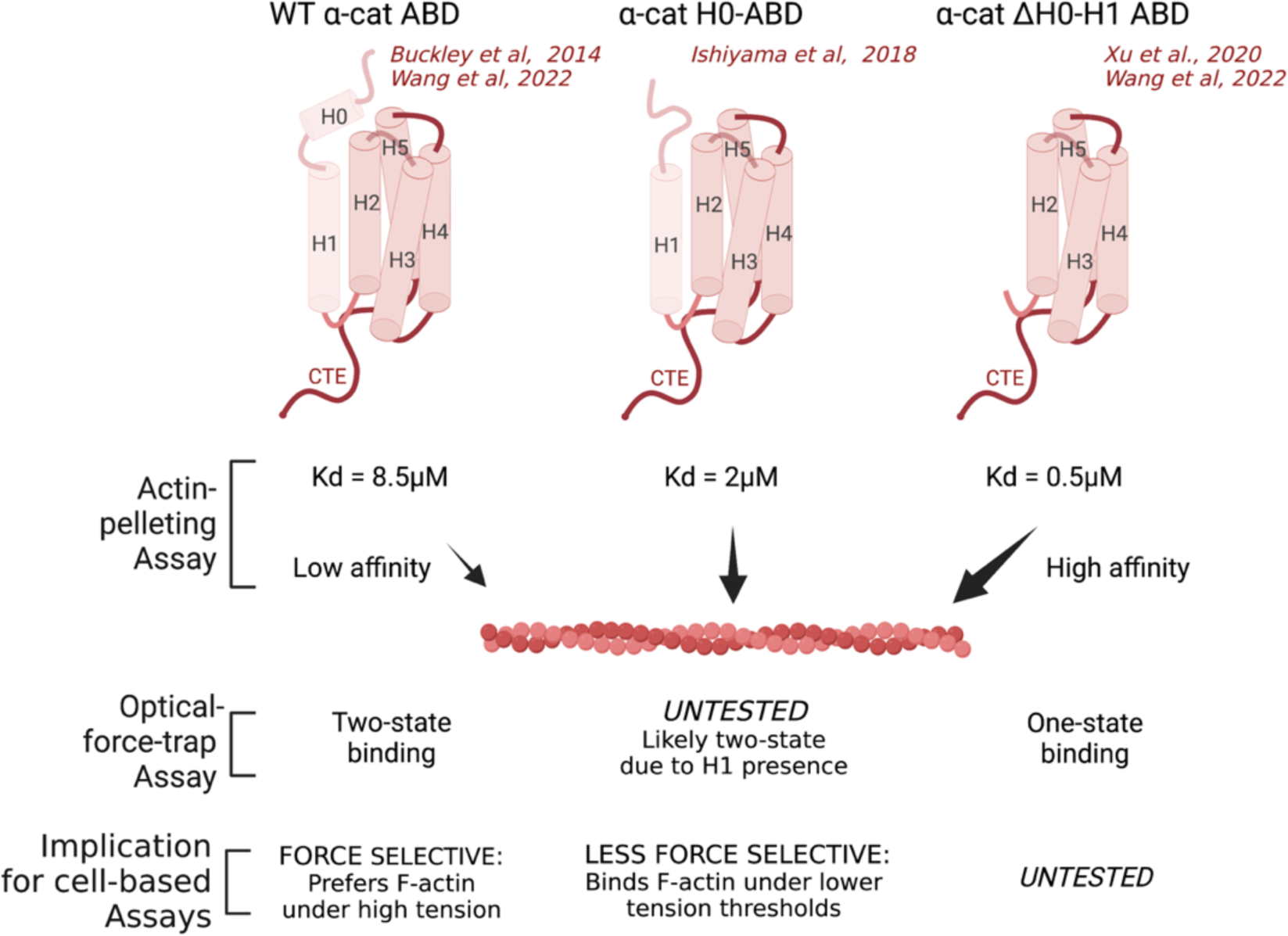
Schematic of α-cat F-actin-binding domain mutants and properties used in this study. (**Left**) Schematic of WT α-cat actin-binding domain based on crystallographic structure, which reveals bundle of 5 helices and an unstructured C-terminal extension (CTE) (Ishiyama et al., 2018; Ishiyama et al., 2013; Rangarajan and Izard, 2012; Rangarajan and Izard, 2013). WT α-cat shows low affinity binding to F-actin in solution (Actin-pelleting Assay, Kd = 8.5μM) and two-state binding by single molecule force-trap measurements (Buckley et al., 2014; Wang et al., 2022; Xu et al., 2020). (**Middle**) Schematic of α-cat-H0-FABD protein designed by Ishiyama et al (2018) to unfold the short H0-helix. This mutant shows enhanced binding to F-actin in solution (Kd = 2μM). Single molecule force-trap measurements remain untested for this mutant; two-state binding is likely possible based similarity to vinculin. (**Right**) The α-cat-ΔH0-H1-FABD protein of Wang et al (2022) removes both H0-H1 helices. This mutant shows dramatically enhanced (18-fold) binding to F-actin in solution (Kd = 0.5μM) and one-state slip-binding by single molecule force-trap measurements (Wang et al., 2022). Cell based studies carried out here (MDCK cells) and elsewhere (colon carcinoma cell line, Ishiyama, 2018 #1555) suggest the α-cat-H0-FABD mutant is less force-“discriminating” or “selective” of actin filaments than WT α-cat protein, with implications for static versus dynamic cell-cell behaviors. Created with BioRender.

